# A thermoplastic chip for correlative assays combining screening and high-resolution imaging of immune cell responses

**DOI:** 10.1101/2024.10.02.616277

**Authors:** Hanna van Ooijen, Quentin Verron, Hanqing Zhang, Patrick A. Sandoz, Thomas W. Frisk, Valentina Carannante, Karl Olofsson, Arnika K. Wagner, Niklas Sandström, Björn Önfelt

**Affiliations:** Department of Applied Physics, Science for Life Laboratory, KTH Royal Institute of Technology, Stockholm, Sweden; Department of Microbiology, Tumor and Cell Biology, Karolinska Institutet, Stockholm, Sweden; Department of Medicine, Center for Hematology and Regenerative Medicine, Huddinge, Karolinska Institutet, Stockholm, Sweden; Haematology Center, Karolinska University Hospital, Stockholm, Sweden; Department of Medicine, Center for Infectious Medicine, Huddinge, Karolinska Institutet, Stockholm, Sweden

**Keywords:** Microwell, Correlative imaging, Screening, High-resolution, Single-cell, 3D cell culture, Spheroid, Tumor microenvironment, Natural killer cell, Serial killing

## Abstract

Single-cell immune assays are developed for the identification and characterization of individual immune cell responses. Some methods provide snapshots of the phenotype of the cell, such as flow cytometry and single-cell RNA sequencing, whereas others, almost exclusively microscopy-based, can be used for longitudinal studies of individual cells. However, obtaining correlative data on cell dynamics and phenotype of individual immune cells is challenging but can provide more nuanced information of heterogeneous immune cell responses. In this work, we have addressed this challenge by developing an easy-to-use, disposable, thermoplastic microwell chip, designed to support screening and high-resolution imaging of single-cell behavior in two-and three-dimensional cell cultures. We show that the chip has excellent optical properties and we provide simple protocols for efficient long-term cell culture of suspension and adherent cells, the latter grown either as monolayers or as hundreds of single, uniformly-sized spheroids. We demonstrate the applicability of the system for single-cell analysis by correlating the dynamic cytotoxic response of single immune cells grown under different metabolic conditions to their intracellular cytolytic load at the end of the assay. Additionally, we illustrate highly multiplex cytotoxicity screening of tumor spheroids in the chip, comparing the effect of environment cues characteristic of the tumor microenvironment on natural killer (NK) cell-induced killing. Following the functional screening, we perform high-resolution 3D immunofluorescent imaging of infiltrating NK cells within the spheroid volumes.

## Introduction

The immune system consists of highly diverse, specialized cells that collaboratively contribute to protecting the host against disease (1). Many of these cells, for example cytotoxic T cells, are restricted to the recognition of a single antigen, meaning that prior to expansion, merely a small fraction of all cells responds against a specific signal (2). As a result of this individualized response, a variety of assays that enable the study of single cells have been applied for immune cell investigations. These include limiting dilution analysis where clones of immune cells are created (3), the detection of secreted molecules using ELISPOT (4,5), flow and mass cytometry (6,7) and single-cell RNA sequencing (8,9). Although informative of single-cell characteristics, most of these assays do not allow detailed longitudinal studies of individual cells, thereby preventing their use for the study of dynamic behavior over extended periods of time, such as contact formation dynamics, migratory potential or serial-killing capacity. For such applications, the spatial separation of single cells using microwells or microfluidics is an attractive alternative (10–15). The isolated cells can then be monitored through time-lapse microscopy, making it possible to dissect transient behavioral patterns and relate cell responses happening on different time scales.

We have previously developed methods to study single-cell responses of natural killer (NK) cells (16– 18). NK cells are innate immune cells capable of rejecting tumor-transformed and virally-infected cells, and to act as immunoregulators of both the innate and adaptive immune systems by secreting cytokines (19). Due to these properties, NK cells have been identified as a potent target for cancer immunotherapy (20). For example, effective treatment of B-cell malignancies with the monoclonal antibody Rituximab has been shown to partially work through antibody-dependent cellular cytotoxicity (ADCC) of antibody-coated tumor cells by CD16-expressing NK cells (21). As the interest in harnessing the anti-tumor potential of NK cells is rising, an improved understanding of the mechanistic features that determine NK cell cytotoxicity is needed. For example, the best clinical effects of NK cell therapy have so far been shown in hematological cancers and questions remain as to why NK cells show dysfunctionality in solid tumors (22,23).

To study the response of immune cells to solid tumors, there is a strong drive to develop *in vitro* models that resemble *in vivo* tumors (24–27). Three-dimensional (3D) tumor spheroids (28,29) are multicellular systems that mimic several key factors important for the tumor microenvironment (TME), such as sustained cell-to-cell interactions and differential nutrient accessibility (30,31). Such models enable the study of how immune cells infiltrate and kill in the 3D environment (32,33).

Tumor spheroids are typically monitored using microscopy-based techniques, yet the structure and size of the spheroids cause light scattering, thereby making highly-resolved imaging of cells in the central and upper parts of the spheroid challenging (34). Consequently, most 3D cell culture techniques only examine tumor spheroids using low-resolution microscopy (34,35). During recent years, new methods to reduce light scattering through tissue clearing have been developed, potentiating the complete scanning of thick specimens with high-resolution microscopy (36). However, most 3D culture protocols involve substrates that are not compatible with high resolution imaging (34,37). To solve this, some approaches involve the transfer or direct creation of tumors in conventional cell culture dishes with a flat, low-adhesion bottom surface. However, once seeded in a large cell-culture dish, the tumors cannot be kept separated over prolonged cultivation times, due to cell migration and mixing during medium exchange, thereby reducing the potential for longitudinal measurements (38). Using our microwell system, we have previously developed an ultrasound-based approach for creating hundreds of tumor spheroids of uniform size, all kept separated during cell culture, in a microscopy-compatible silicon chip with a #1.5H glass bottom (39–41). However, silicon-glass chips are relatively expensive compared to other lab consumables like various plastic cell culture substrates and the method requires an external ultrasound generator to create tumor spheroids.

In this work, we present a methodology for longitudinal, high-content screening and subsequent high-resolution imaging of individual immune cells in both 2D and 3D cell cultures. For this purpose, we developed novel plastic microwell chips, which are transformative in that they combine the merits of conventional lab consumables, like microplates, with the performance of specialized research technology, like silicon-glass microwell chips. These new disposable microwell chips are evaluated for their optical quality, liquid and cell handling, and for supporting various cell cultures, which include highly parallelized formation of multicellular spheroids directly in the flat-bottom microwells. Here we exemplify the methodology by applying it to study NK responses in 2D and 3D tumor cell cultures under multiple conditions that simulate the TME. Time-lapse microscopy is combined with subsequent high-resolution analysis of the immune and tumor cell co-cultures in the chip. Our results confirm the potential of the microwell chip-based methodology to enable a wide range of correlative single-cell analysis involving both static and longitudinal parameters.

## Results

### The plastic microwell chips have optical quality compatible with super-resolution microscopy

We developed injection molded thermoplastic microwell chips to enable high-content screening with subsequent high-resolution imaging of individual cells in an accessible and microscopy-oriented format. The chips are available in two versions. One version contains 4 chambers, with a volume of 438 µL each; the second, 16 chambers with a volume of 105 µL (Fig. 1A). All chambers include a microwell array at the bottom, containing 169 or 36 microwells, respectively, with a volume of 48 nL each. In both designs, the well diameter and pitch were optimized so that 16 wells fit within the imaging field-of-view (FoV) using a microscope with a 5x objective and a camera sensor size of 13×13 mm^2^; four wells when using a 10x objective; or one well within the FoV of a 20x objective (Fig. 1A, Supp. Fig. S1A-C). For an easy set-up on the microscope and an efficient screening procedure, all imaging positions in the chip can be accessed by an integer number of steps from any corner. The 450 μm depth of the wells allows liquid to be removed from the chambers without emptying or even creating a substantial fluid flow inside the wells, thereby avoiding cell disturbance during medium exchange (Fig. 1B). Alternatively, the cells can be retrieved from the wells by vertical pipetting directly above the wells, then collecting the cell suspension from the entire chamber (Fig. 1B). We demonstrated these capabilities in a chip containing fluorescent beads by first performing 10 successive complete liquid exchanges using the low-angle pipetting technique, followed by full bead retrieval using the vertical pipetting technique (Supp. Movie 1). In addition to the chips, we designed both petri dish-format and microplate format chip holders, fitting in standard microscope stage inserts (Fig. 1C). The petri dish-format holder has a single chip slot whereas the microplate-format holder has 8 slots, thereby extending the number of conditions that can be used in a single experiment from 16 up to 128. The design of the chip holder can be adapted to any of the standard substrate formats used in microscopy, making the integration of the microwell chips to various assays and workflows flexible. A plastic lid was also designed and produced by injection molding. The lid covers the chip to limit evaporation and contamination, while still allowing gas access to the chambers. As an alternative solution, the top edge of the 4-chamber microwell chip contains ridges intended to support a standard 22×22 mm^2^ coverslip with maintained gas exchange. Photographs show the fabricated chips and lid (Fig. 1D) mounted in holders (Fig. 1E-F). Brightfield microscopy images of individual microwells from each type of chip (Fig. 1G-H), as well as cross-section images (Fig. 1I-J), show flat and smooth bottom surfaces of the produced microwells.

**Figure 1.**
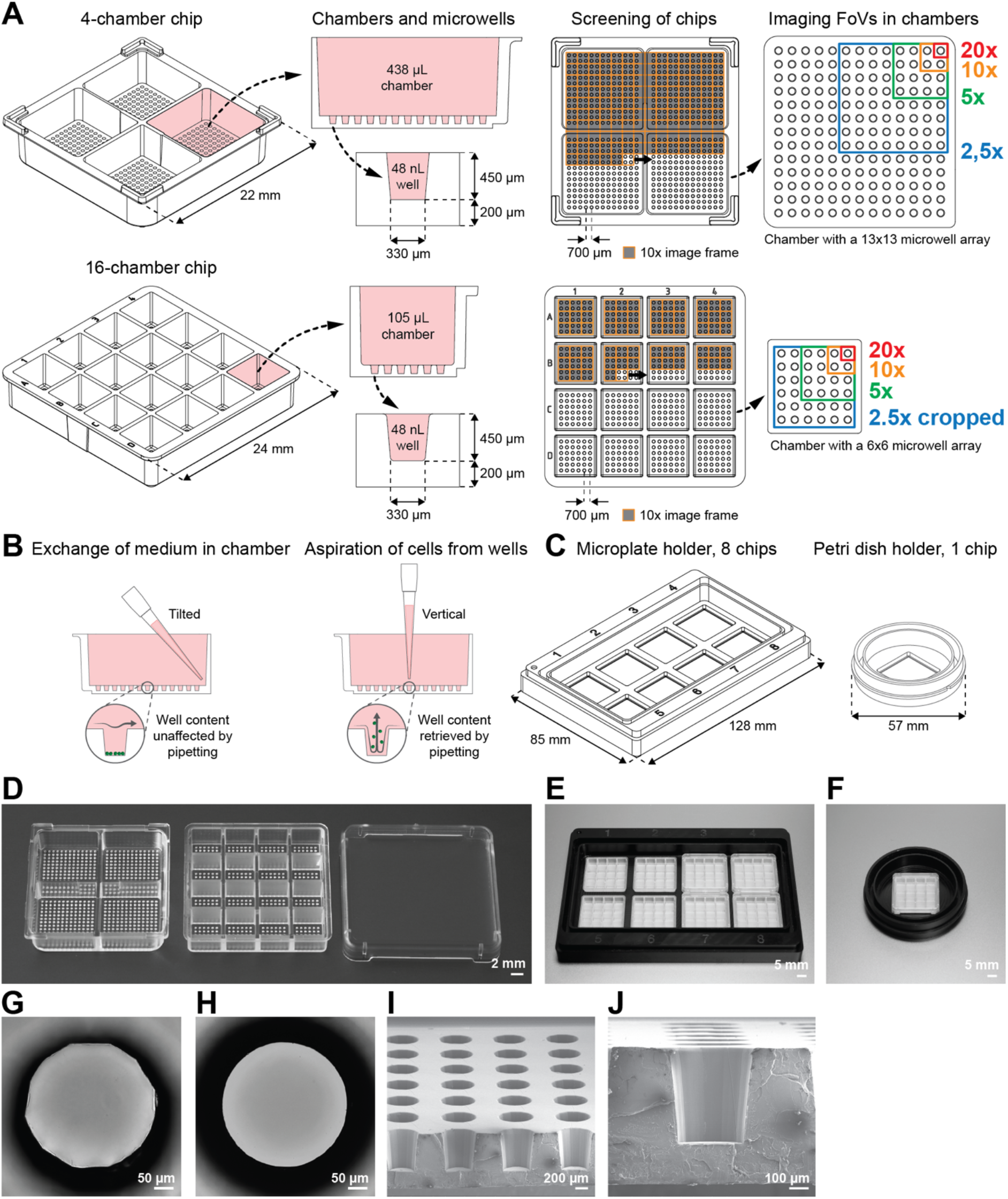
Multichambered plastic microwell chips for 2D and 3D imaging-based cell assays and analysis. **(A)** Illustrations of the 4-chamber (top) and 16-chamber (bottom) microwell chips with cross-sectional views of chambers and microwells as well as top views of the frame-by-frame microscopy screening for each chip and the relative size of the imaging field-of-views at different magnifications. **(B)** Cross-sectional illustrations of a chamber of the 4-chamber chip, showing liquid exchange by tilted pipetting in the chamber without disturbing the content of the wells (left) and cell retrieval from the microwells at the bottom of the chamber by vertical pipetting (right). **(C)** Illustrations of a microplate-format chip holder with eight chip slots (left) and a petri dish-format chip holder with a single chip slot (right). **(D)** Photograph of the produced 4- and 16-chamber plastic microwell chips and the accompanying lid. **(E-F)** Photographs of the produced 8-slot microplate-format chip holder (E), and the 1-slot petri dish-format chip holder (F). **(G-H)** Microscopy brightfield images of a single microwell in the 4-chamber chip (G), and in the more refined 16-chamber chip (H). **(I-J)** Scanning electron micrographs of the cross-section of a microwell array in the bottom of a chamber of a split 4-chamber chip (I), and a close-up of a single microwell (J).

The chips were designed with the intended use of high-content, high-resolution, microscopy-based analysis of live cells. Common objectives are optimized for #1.5 (0.17 mm +-10%) glass coverslips, thereby making resolution highly dependent on the substrate thickness and refractive index. Therefore, the chips were produced with a well-bottom thickness close to that of a glass coverslip #1.5 (with a final thickness of 0.2 mm), using three different thermoplastic materials: polystyrene (PS), cyclic olefin polymer (COP) and cyclic olefin copolymer (COC), all previously proven to be biocompatible (42,43) and used for microfabrication (44). COP and COC were chosen as they have refractive indices close to that of glass (1.530 and 1.533 compared to 1.52 for glass), and PS (refractive index of 1.59) was used as a comparison as it is a common substrate in commercial cell culture vessels. Additionally, to investigate the effect of the well bottom thickness on the optical properties of the chips, we produced chips with three additional bottom thicknesses, 0.22, 0.25 and 0.3 mm, in COC.

To evaluate the optical performance of the chips, we first examined the lateral and axial fluorescence distribution of 500 nm-wide fluorescent beads (Fig. 2A). As a reference, we included a glass coverslip of thickness #1.5 and a standard plastic 96-well plate. The 1 mm-thick bottom of the 96-well plate imposed the use of a long-working distance 40x objective for the comparison. All chip materials showed performance similar to that of glass, and superior to that of the culture plate, both laterally and axially (Fig. 2B-C). Using a long-working distance objective came at the price of a low numerical aperture (NA) and thereby limited resolution, which was not sufficient to rank the optical performance of the chips. For this reason, further comparison was performed using 100 nm-diameter fluorescent beads with an oil-immersion 63x objective, and the 96-well plate had to be excluded from the comparison due to its incompatibility with such objectives (Fig. 2D-E). Lateral resolution was still comparable between all materials tested (Fig. 2D). Axially, COC and COP performed comparably, whereas PS demonstrated reduced resolution, possibly due to the larger refractive index-mismatch between PS and the commercial immersion oil (Fig. 2E). Increasing the well bottom thickness did not affect the lateral resolution, whereas the axial resolution was reduced with thicker materials (Supp. Fig. S2A-B).

**Figure 2.**
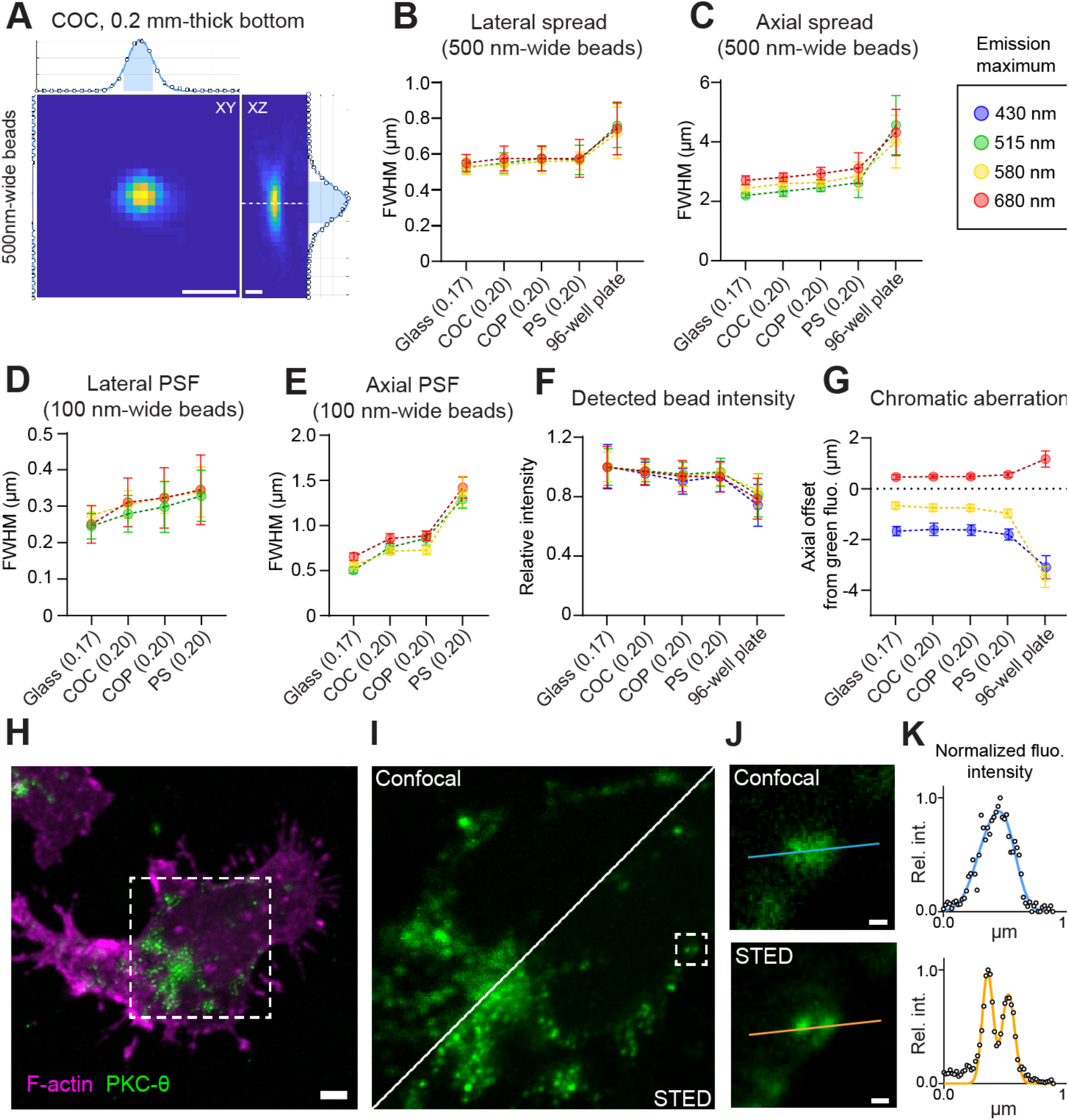
Characterization of the optical performance of the plastic microwell chips. **(A-G)** Fluorescent microbeads were seeded in the wells of various materials and bottom thicknesses and imaged by confocal microscopy to estimate the optical performance through the different chip substrates. **(A-C)** Example top- and side-view of 500 nm-wide beads acquired using a long-distance 40x/0.60 objective through a COC chip with 200 µm-thick bottom (A). Width of the lateral (B) and axial (C) PSF of 500 nm-wide beads. n=18-78 beads per condition. **(D-E)** Sub-diffraction 100 nm-wide beads seeded in the chip microwells were imaged with an oil-immersion 63x objective. Width of the lateral (D) and axial (E) PSF. Pooled data from 3 independent experiments, with a total of n=94-129 beads per condition. **(F)** Fluorescence intensity of 4 µm-wide beads imaged with an 10x objective, normalized to the mean intensity of the beads through a glass coverslip. n=23-98 beads per condition. **(G)** Chromatic aberration through the microwell chips, measured as the axial offset of each fluorescence channel from the green (488 nm) channel, for 500 nm-wide beads imaged with a long-working distance 40x objective. n=15-87 beads per condition. **(H-K)** NK cells imaged by STED microscopy in microchips coated with antibodies against the NK cell receptors CD16 and LFA-1. Example fluorescence image of filamentous actin and the kinase PKC-Θ, labeled in primary human NK cells building immune synapses with the well surface of a COC chip with 200 µm-thick bottom (H). Diffraction-limited PKC-Θ microclusters were further imaged by STED microscopy (I). Example of signaling clusters only resolved in the STED image (J). Fluorescence intensity profiles along the lines indicated in J (K). Scale bars: (A): 500 nm; (H): 1 µm; (J): 100 nm.

Besides resolution, we also investigated the transparency of the chips by measuring the transmitted light intensity of fluorescent beads, emitting at four different wavelengths (Fig. 2F). All chips performed well, with intensities greater than 90% of that measured through glass, a sharp contrast to the intensities seen using a PS culture plate, which exhibited a 25% intensity loss (Fig. 2F). As commercial objectives are typically designed to be used with #1.5 glass coverslips, optical aberrations may appear if imaging is instead conducted through a layer of a different material or thickness. Due to this, we investigated if chromatic aberrations were being induced in our chips, by measuring the axial distance between focal planes for rays of different wavelengths (Fig. 2G). COC and COP chips performed very similarly to glass, however, PS showed a larger axial spread between wavelengths (Fig. 2G). Here again, the 96-well plate exhibited the worst performance. Across all our measurements, the chips manufactured in COC with the thinnest bottom (200 µm) performed best.

As the COC chips with thinnest microwell bottom demonstrated optical properties very close to that of a glass coverslip, we assessed whether this chip could support super-resolution microscopy of biological samples. We performed stimulated emission-depletion (STED) microscopy of diffraction-limited signaling micro-clusters in NK cells triggered to interact with the chip surface (Fig. 2H-J). Using STED, we gained an improved resolution with single spot lateral PSFs as narrow as 130 nm compared to over 250 nm with confocal microscopy (Fig. 2D, K). Thus, the COC chip with a 200 µm-thick well bottom allowed super-resolution microscopy, offering axial and lateral resolutions compatible with the study of intracellular micro-structures with sufficient optical sectioning. Taken together, the optical characterization demonstrated that the COC chip with 200 µm bottom thickness performed almost as well as #1.5 glass coverslips making it our chip design of preference for the rest of the study.

### Plasma-treated COC chips support two-dimensional cell culture in microwells

The microwell chips were primarily designed for microscopy-based screening of cell function, and therefore, we confirmed that the platform was compatible with long-term cell culture. As untreated COC plastic does not promote cell adhesion (45), the plastic chips were treated with a low-pressure oxygen plasma and the resulting charge on the plastic surface efficiently supported cell attachment (Fig. 3A). Five commonly used adherent human tumor cell lines, ovarian cancer OVCAR-8, colon cancer DLD-1, cervical cancer HeLa, breast cancer MCF-7 and neuroblastoma IMR-32, and five suspension human cell lines, chronic myelogenous leukemia K562, acute T cell leukemia Jurkat, acute lymphocytic leukemia Nalm-6, acute lymphocytic leukemia SUP-B15 and acute myeloblastic leukemia Kasumi-1, were successfully grown in the chip (Fig. 3A). All cell lines divided within 48 hours, and the cell division rates were similar to those reported by cell line repositories ATCC and DSMZ (Supp. Fig. S3A-B).

**Figure 3.**
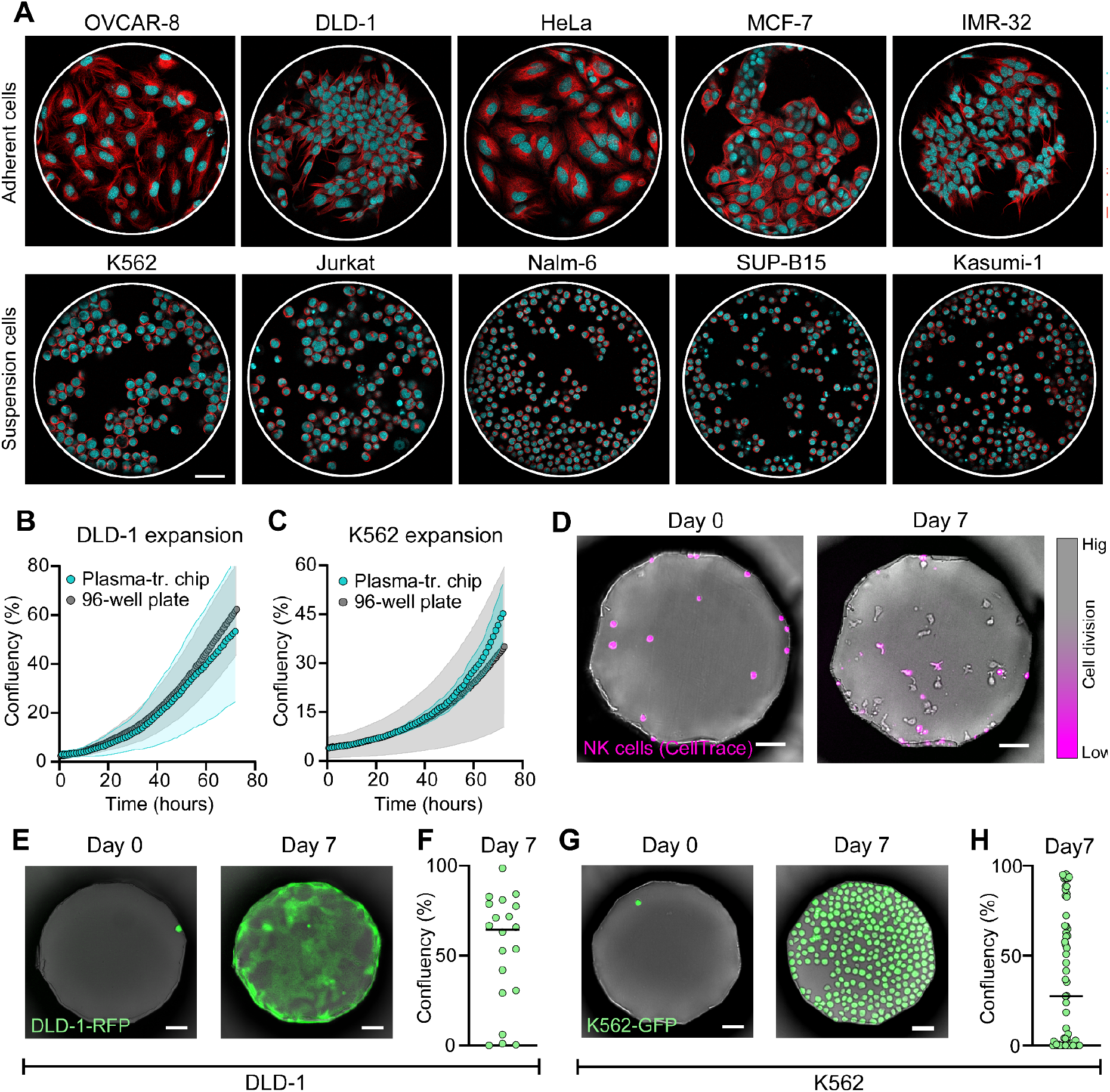
Compatibility of the microwell chip with suspension and adherent 2D cell culture. **(A)** Representative fluorescence microscopy images of adherent and suspension cell lines growing in the wells of plasma-treated chips after 48 hours of culture. **(B-C)** Cell growth in the plastic chips was compared to that in a standard 96-well plate. Average confluency over time of the adherent cell line DLD-1-RFP (B) measured in 30 wells in the chip and in 21 imaging positions in the 96 well plate, and of the suspension cell K562-GFP (C) measured in 13 wells in the chip and 16 imaging positions in the 96 well plate. The wells were selected so that the average starting cell density was similar for both conditions. **(D)** Example images of CellTrace-labeled primary human NK cells grown in the microwell chip. **(E)** Example images of a clonal expansion generated from a single DLD-1-RFP cell after 7 days of culture. **(F)** Cell confluency obtained after 7 days in wells containing a single DLD-1-RFP cell at seeding. Each dot represents a single well, n=20. **(G)** Example images of a clonal expansion generated from a single K562-GFP cell after 7 days of culture. **(H)** Cell confluency obtained after 7 days in wells containing a single K562-GFP cell at seeding. Each dot represents a single well, n=59. Scale bars: 50 µm. The error bars in (B-C) denote standard deviation, and the black lines in (F,H) the median value.

The kinetics of cell division were further investigated using one adherent cell line, DLD-1, and one suspension cell line, K562, stably transfected to express a fluorescent protein (DLD-1-RFP and K562-GFP). The growth rates were similar for cells grown on a plasma-treated COC chip and on a standard, cell culture-treated, polystyrene 96-well plate (Fig. 3B-C). Unlike the adherent cells, the suspension cell line K562 was also capable of growing equally well on untreated plastic chips (Supp. Fig. S3C). We also tested whether primary human cell culture could be performed in the chip. Peripheral blood mononuclear cells (PBMCs) and purified human NK cells were efficiently cultured and expanded in the chip for 7 days (Fig. 3D, Supp. Fig. S3D-E). Cell expansion was seen also when the cells were seeded at very low cell densities.

As the chip confines individual cells to their respective wells, we investigated whether we could utilize the microwell array to generate clonal expansions of single cells. To do so, we seeded DLD-1-RFP or K562-GFP cells at low concentration and monitored wells in which a single tumor cell was originally seeded (Fig. 3E-H). A complete media exchange was performed every two days without disturbing the cells, thanks to the limited liquid flow inside the deep wells. Most cells managed to form single cell expansions (Fig. 3F,H), although the number of cell divisions reached after 7 days was highly variable between single cells. Such division heterogeneity has been reported previously, even for single cells isolated from the same cell line (46–48). We further investigated whether the chip itself had any influence on cell division by comparing cell proliferation between chambers, between different regions inside a chamber, and between different cell seeding densities, but no differences were observed (Supp. Fig. S4A-D). These experiments illustrate how the chip can be used to study the growth behavior of individual cells, including non-adherent cells (Fig. 3G-H). Altogether, we could demonstrate that the plasma-treated microwell chip allowed efficient 2D cell culture of both tumor cell lines and primary immune cells, even when starting from single cells.

### The microwell chips facilitate the investigation of single NK cell function

We decided to apply the microwell chips to study immune cell function in cancer, and more specifically, the effect of the TME on NK cell function. Activated NK cells are highly motile immune cells, and therefore require a short duration between time-lapse frames to be successfully tracked during single-cell functional evaluation. The 4-chamber microwell chip contains 169 wells per chamber, and typically, we find that at least 6 highly motile cells can be analyzed per well, using an experimental design with 4 fluorescent markers and an imaging time resolution of 3 minutes. This results in a total of approximately 1000 analyzable cells per each of four conditions. These relatively high cell numbers make the 4-chamber chip design particularly useful for the study of functional properties of rare cell populations. To demonstrate the potential of the microwell chip for single-cell tracking and functional assessment, we created co-cultures of the lung cancer cell line A549 and primary NK cells. Prior to the co-culture, we treated NK cells in four different environments, chosen to simulate conditions met in the TME; low glucose level, glutamine restriction, high lactate level, along with an untreated control. After 24 hours of treatment, the NK cells were added to A549 cells grown in a 4-chamber chip. The chip was then imaged every 3 minutes for 5 hours (Fig. 4A). Similarly to other microscopy-based killing assays such as the commercial Incucyte assay (Sartorius), we generated bulk killing curves of the population average kinetics of NK cell killing (Fig. 4B) (49). Here, dead cells were identified automatically, either using a fluorescent dead cell marker and the open-source image analysis platform CellProfiler, or only the bright-field channel and a machine learning-based detection method developed in-house (Fig. 4B, Supp. Movie 2). Although informative of differences in the total killing levels, as well as the kinetics of the killing, these bulk measurements do not provide insight into the proportion of active NK cells, or how the killing capacity of individual cytotoxic cells varies. Thus, we made use of the highly time-resolved data, and tracked individual NK cells in order to classify their killing ability (Fig. 4C-D). The majority of NK cells displayed cytotoxic potential, however all TME treatments reduced the fraction of cytotoxic cells (Fig. 4E). In all four conditions, we observed a small group of cells (∼15 % of all NK cells in the control condition) that were particularly efficient killers, killing 3 or more targets (Fig. 4F). Interestingly, although only small differences in the fraction of cytotoxic cells could be observed when comparing the three TME treatments, the fraction of serial-killing NK cells was largely affected by glutamine restriction, suggesting that serial killing might be dependent upon distinct metabolic requirements (50).

**Figure 4.**
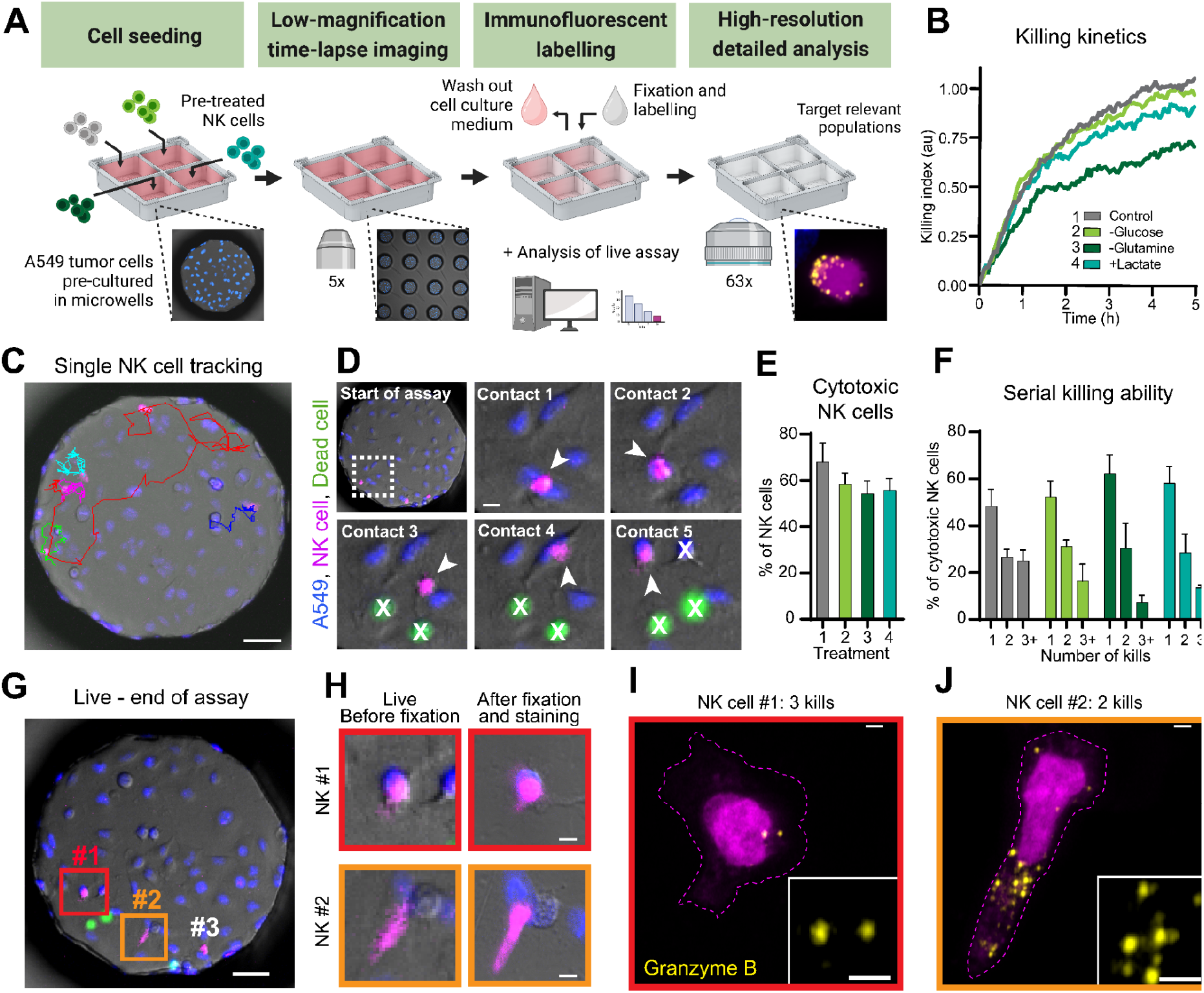
Application of the microwell chip to study the reduced NK cell anti-tumoral function under TME conditions. **(A)** Illustration of a microwell chip-based killing assay using primary NK cells and an adherent target cell line, involving a live, time-lapse cytotoxicity assay of NK cells, followed by fixation and immunofluorescence labeling of the co-cultures. **(B)** Bulk killing kinetics of A549 cells by NK cells, measured as the number of dead target cells in each time point, divided by the number of NK cells in the well at the start of the assay (killing index). Each curve represents the mean from 3 independent experiments using 3 biological replicates, with the result of each experiment being calculated as the mean of at least 96 wells. The wells were selected so that the average starting NK cell density was similar and target cells were always in excess for all conditions. **(C-F)** Single NK cells were followed over time during their interaction with A549 tumor cells to quantify single cell killing. Example image of migration tracks from 5 individual NK cells during the 5h killing assay (C). Sequence of contacts formed by a serial killing NK cell during the assay (D). Arrowhead: NK cell in contact; X: dead target cell. Scale bar: 10 µm. Proportion of cytotoxic NK cells (E). Fraction of the cytotoxic NK cells killing 1, 2 or 3 or more target cells during the assay (F). The bars in (E-F) represent the mean and standard deviation of 3 independent experiments using 3 biological replicates. A minimum of 68 cells were analyzed for each condition. **(G-J)** At the end of the time-lapse assay, the co-cultures were fixed and the NK cells were fluorescently labeled for the cytolytic molecule granzyme B. Overview of a co-culture well at the end of the assay (G). Scale bar: 50 µm. Example images of NK cells imaged at the end of the time-lapse assay (left) and after fixation (right) (H). Scale bars: 5 µm. Example immunofluorescence images of the granzyme B content in a serial killing NK cell (I) and a less cytotoxic NK cell (J), after having interacted with A549 cells. Scale bar: 1.5 µm.

As discussed earlier, the geometry of the microwells allows culture media to be exchanged in the chambers, without disturbing the cells in the wells. Thanks to this property, we could fix and label the cells in the wells after the end of the cytotoxicity assay (Fig. 4G-H). As the cells stay in place during this process, we could directly correlate the dynamic cytotoxic response of individual cells to intracellular markers, including those requiring high-or super-resolution imaging, such as in this case, granzyme B expression (Fig. 4H-J, Supp. Movie 3). These experiments demonstrate the potential of the plastic microwell chip for single-cell studies, even of highly motile suspension cells. Being able to fully exchange the medium in the chambers without altering the cell positions opens the possibility for a variety of new correlative assays.

### Straightforward tumor spheroid formation with maintained optical access

Traditionally, cellular assays have been carried out using 2D co-cultures; however, when studying adherent target cells, questions arise as to whether the observed results are biologically relevant. For this reason, we developed and evaluated protocols that allow 3D cell culture in the plastic microwell chips (Fig. 5A). The well geometry and the low cell adherence on the surface of untreated COC allowed adherent DLD-1 cells to spontaneously form tumor spheroids in the wells (Supp. Fig. S5A). However, these spheroids often showed partial adherence to the chip surfaces, resulting in uneven tumor shapes and focus problems as cell aggregates attached to the well walls (Supp. Fig. S5A). To avoid this, the chip was coated with an anti-adherence solution which prohibited cells in spheroids from binding to the well surface (Supp. Fig. S5B). We compared the dynamics of suspension cell growth and tumor spheroid formation on non-coated and anti-adherence-coated chips and found similar cell growth rates and tumor distributions, demonstrating that the coating solution was well tolerated (Supp. Fig S5C-D). Cell aggregates coming in contact during the assay typically merged and rounded up into a single, larger aggregate. However, many wells still contained several tumor spheroids after 72h hours of culture as the clusters remained separated in the well. Therefore, we developed a protocol to encourage cell-to-cell contacts by tilting the chip by approximately 15 degrees for 24 hours following cell seeding (Fig. 5A). After this, the chip was placed in a horizontal position again and microscopy could be performed. Tilting the chip greatly increased the proportion of wells with a single tumor spheroid per well after 48 hours of culture (Fig. 5B). Combining anti-adherence treatment with tilting of the chip, we could produce uniformly sized, single spheroids in close to all 676 microwells on the chip (Fig. 5B-C, Supp. Fig. S5E-F).

**Figure 5.**
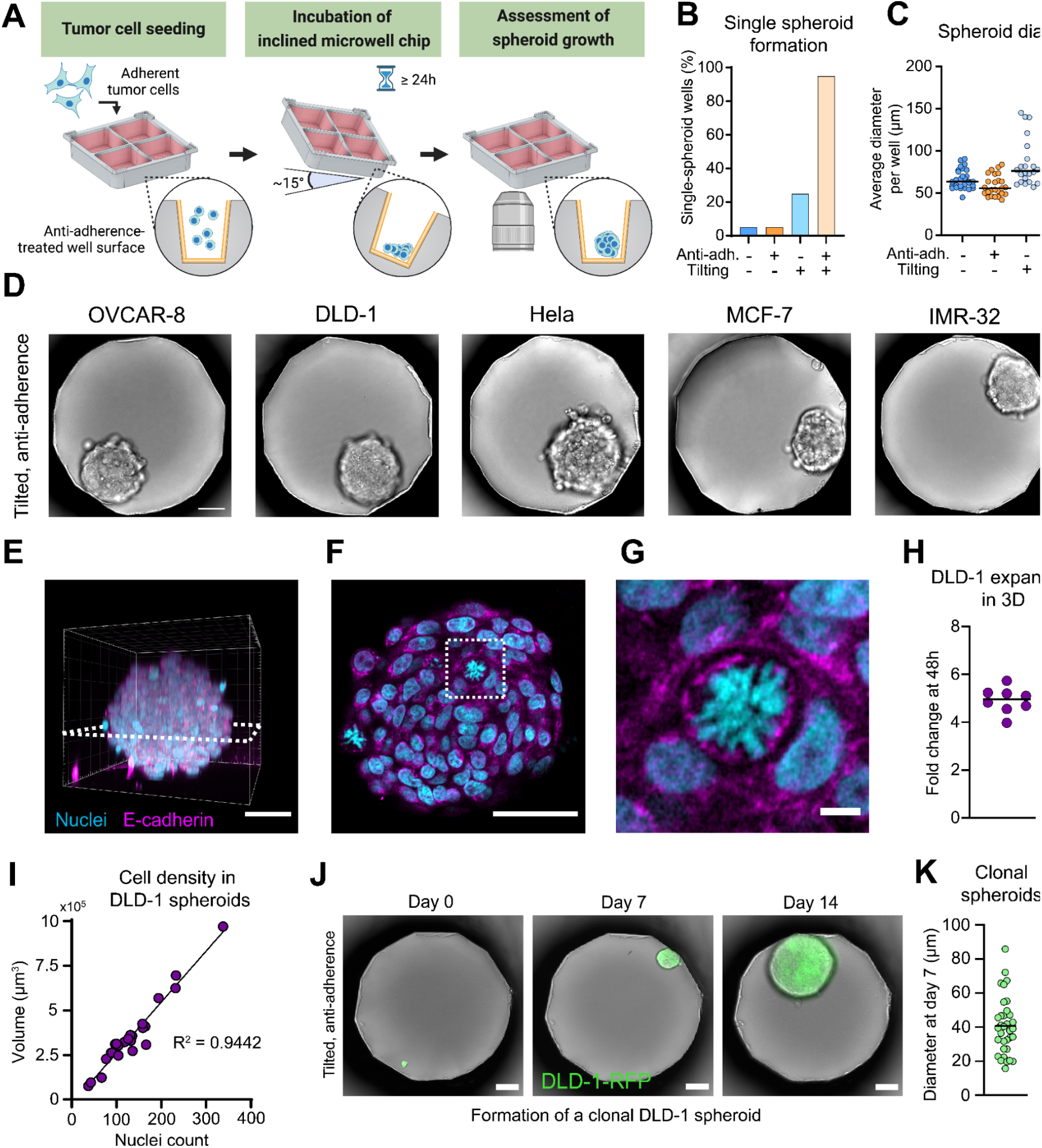
Optimization of a 3D cell culture protocol for spheroid formation in the microwells. **(A)** Schematic illustration of the spheroid formation workflow. Adherent cells are seeded in the anti-adherence-treated microwells, and kept in culture for 24 hours in the chip, tilted at 15 degrees to facilitate intercellular contacts. **(B)** Fraction of wells containing a single tumor spheroid after 48 hours of culture. n=20 wells/condition. **(C)** Median diameter of tumor spheroids in the wells after 48 hours of culture. Each dot represents the spheroid diameter in a single well. n=22-33 wells/condition. **(D)** Representative bright-field images of 3D spheroids of 5 different cell lines, after 48 hours of culture in tilted, anti-adherence-treated chips. **(E-G)** 3D reconstruction (E) and single-plane (F-G) confocal images of a fluorescently-labeled, optically-cleared DLD-1 spheroid in a microwell. **(H)** Growth of DLD-1 spheroids in the wells, expressed as an average fold change after 48 hours from the cell count in the well at seeding. Each dot represents a single well, n=8. **(I)** Correlation between the spheroid volume and the number of segmented nuclei, for DLD-1 spheroids grown for 48 hours. The line represents a linear regression. Each dot represents a single spheroid, n=25 spheroids. **(J)** Example wide-field microscopy images of a clonal spheroid growing for 14 days from a single DLD-1-RFP cell. **(K)** Average diameter of the spheroids obtained from single cells after 7 days of culture. Each dot represents the spheroid diameter in a single well. n=29-39 wells/condition. Scale bars in (E,F,J): 50 µm. Scale bar in (G): 5 µm.

Using the optimized protocol, 14 adherent cell lines were tested for their ability to form tumor spheroids in the chip (Fig. 5D). All cell lines except renal ACHN consistently formed tumors (Supp. Table 1). We validated that the tumors were in fact spherical by performing full-volume confocal microscopy of optically cleared tumors (Fig. 5E-F). Clearing was performed as this greatly improves the imaging quality of large spheroids that otherwise suffer from severe light scattering (Fig. 5E-G, Supp. Fig. S6A-B). By automatic 3D segmentation and counting of the stained nuclei, we also measured the growth rate of the DLD-1 cells grown in 3D (Fig. 5H). To investigate if the cell density varied for spheroids of different diameters, we purposely seeded cells unevenly in the microwells, providing a large heterogeneity in the number of cells per well at start, and imaged the chip after 48 hours (Fig. 5I). Interestingly, we found that the spheroid volume was directly proportional to the number of cells, i.e. the cell density was independent of the spheroid size for this cell line.

Since we successfully generated 2D clonal expansions from single cells in the wells, we investigated whether the chip would also allow the formation of entire tumor spheroids from individual cells. We seeded adherent cells DLD-1-RFP at a low density and followed wells that had received a single cell upon seeding (Fig. 5J). Two thirds of all individual cells managed to form spheroids after 7 days of culture, and these cultures could be maintained for at least 2 weeks (Fig. 5J, Supp. Fig. S5G). We observed a large variability in the dimensions of 7 day-old tumor spheroids, again demonstrating that the cell division rates vary between single cells of the same cell line (Fig. 5K). In summary, we developed a straightforward protocol for the on-chip formation of hundreds of uniform spheroids from different cell types. The spheroids could then be analyzed live, or fixed and stained for high-resolution imaging directly in the chip, thus minimizing the risk of artifacts introduced by transferring the spheroids.

### The microwell chip enables correlative functional assays of tumor spheroids

Having established an efficient protocol for 3D cell culture, we proceeded to assess the cytotoxicity of NK cells against 3D microtumors formed by A549 tumor cells. Building upon our experiment with 2D co-cultures, we designed a proof-of-principle experiment where NK cells were pretreated with conditions designed to mimic the TME. As immune cells are commonly exposed to hypoxia in cancer tissues, we additionally included low oxygen concentration as a treatment. Using the 16-chamber chip, we could simultaneously investigate all four treatments (low glucose level, glutamine restriction, high lactate level and low oxygen level), as well as all combinations thereof, along with a negative control (Fig. 6A). After 24 hours of pretreatment, the NK cells were added to A549 microtumors in the wells, and the entire chip was imaged every 15 minutes for 16 hours (Fig. 6B-D). When comparing A549 killing in the chip chambers, it was clear that NK cells showed most sensitivity to combined TME treatments, as tumor cell killing was only slightly reduced in the single treatments (Fig. 6E). Combining all treatments resulted in decreased killing and altered killing dynamics, with NK cell cytotoxicity reaching a plateau sooner than in single treatments, where it was not reached by the end of the 16 hour-long assay (Fig. 6F). Surprisingly, we found that in combination, certain conditions appeared to cancel each other and result in restored or even increased killing, even compared to control. For example, combining any treatment with high lactate resulted in mostly unchanged or slightly improved killing compared to the treatment alone, and combinations that included glutamine and oxygen restriction led to poor killing, even if statistical significance was not reached for this limited dataset.

**Figure 6.**
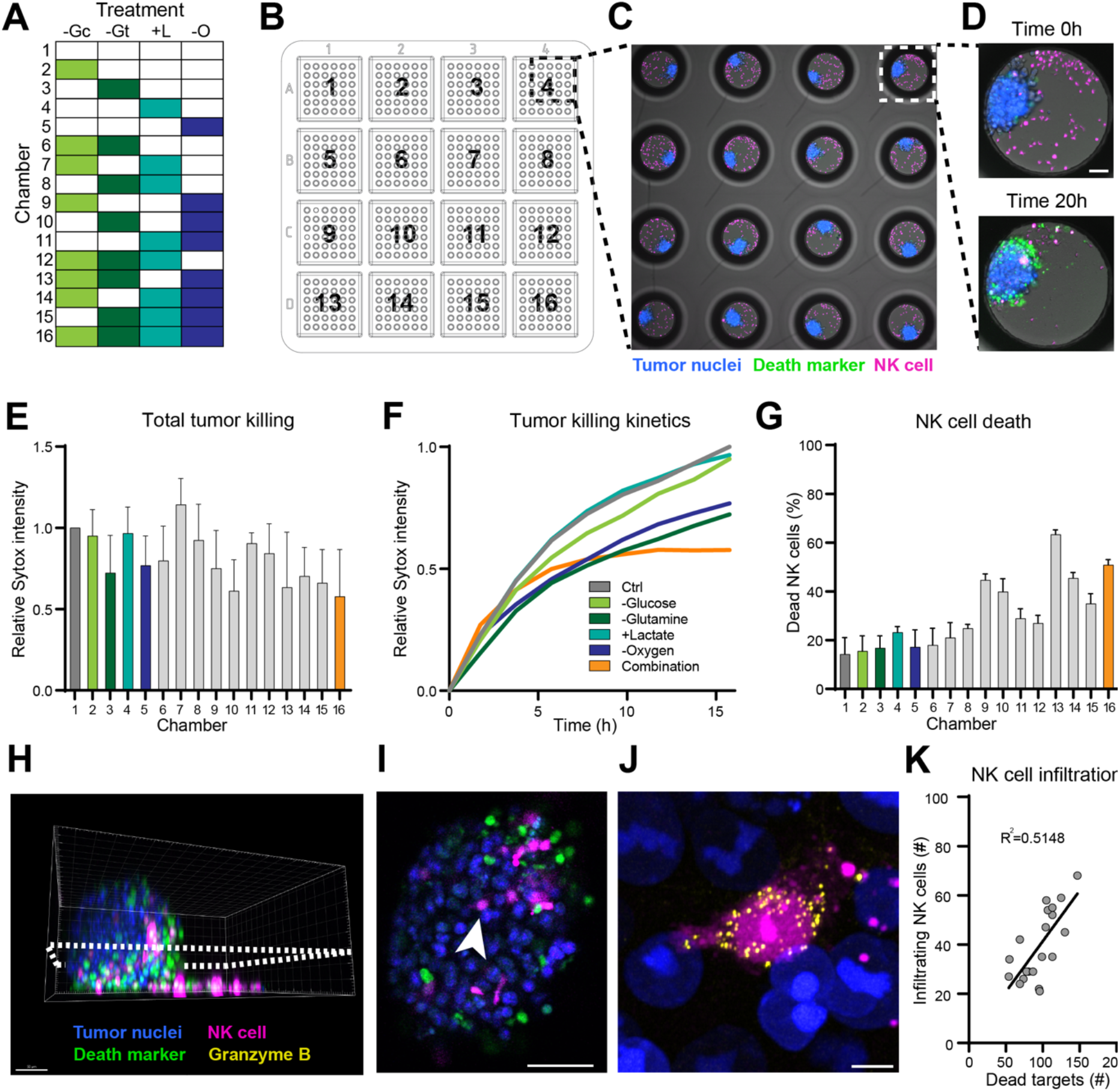
Effect of the combination of TME treatments on NK cell killing of A549 spheroids. **(A)** Primary NK cells were pretreated with 16 conditions designed to mimic the tumor microenvironment, before being challenged against A549 tumor spheroids. -Gc: low glucose level, -Gt: glutamine restriction, +L: high lactate level, and -O: low oxygen level. **(B)** Overview of the experiment layout in the chip, with NK cells from each pretreatment assigned to a plastic chip chamber. **(C)** Example image of the cytotoxicity assay against tumor spheroids. The image is an entire FoV acquired with a 5x objective. **(D)** Example images of the NK cell - tumor cell co-culture in a well, imaged at the beginning (top) and the end (bottom) of the assay. **(E)** Average Sytox intensity per microwell at the end of the 16-hours killing assay, relative to that of the control condition. Each bar represents the mean and standard deviation of 3 independent experiments using 3 biological replicates. **(F)** Killing dynamics, as measured by the relative Sytox intensity over time presented for a subset of the treatments in (E). **(G)** Percentage of dead NK cells at the start of the assay. In (E-F), the same number of live NK cells were added to each chamber at the start of the assay. **(H-J)** Confocal 3D imaging of cleared NK cell - tumor co-cultures at the end of the assay, acquired using low magnification (20x/0.8) for the whole spheroid volume (H-I) and high magnification (63x/1.4) for single infiltrated NK cells (J). Scale bars: 50 µm (H-I), 5 µm (J). **(K)** Number of untreated NK cells on the surface of or inside the spheroids at the end of the killing assay, compared to the number of dead tumor cells detected in the spheroid. The black line represents a linear regression.

The negative effect of both single and combinatorial treatments on NK cells was evident when measuring the proportion of dead NK cells due to the pretreatment (Fig. 6G). Although the same number of live NK cells were added to all chambers of the chip for the killing experiments (Fig. 6E-F), these results indicate that the negative effect of the TME on NK cell activity could be due to an overall reduced NK cell fitness as well as specific effects on their cytotoxic potential.

Similar to the process described for 2D co-cultures, we fixed the co-cultures of NK cells and A549 microtumors at the end of the killing assay, and proceeded with immunofluorescent labeling of granzyme B directly in the wells (Fig. 6H-J). Embedding the co-cultures in a refractive index-matched medium resulted in optically cleared samples, which made it possible to resolve single cytolytic granules of 200 nm size in tumor-infiltrating NK cells by airyscanning microscopy, even deep in the spheroid volume (Fig. 6J, Supp. Movie 4). We also assessed the number of NK cells crawling on or having infiltrated tumor spheroids in the control condition by the end of the assay, and found a strong correlation to the count of dead target cells (Fig. 6K). These results suggest that NK cell cytotoxicity in 3D co-cultures depends on their mobility around and into the microtumors.

## Discussion

In this study, we described a one-time-use, thermoplastic multi-chambered microwell chip designed for single-cell studies in 2D and 3D. While purposely easy and efficient to use, our plastic microwell chips combine the advantages of conventional multiwell plates with miniaturization. Similarly to multiwell plates, the open architecture with multiple chambers allows for easy liquid exchanges by pipetting and the use of many experimental conditions for screening purposes. On the other hand, the miniaturized microwell arrays provide a high level of parallelization, low liquid volumes and a confined environment in which cells can be monitored over time and across different imaging platforms. These features are shared with microwell chips made in other materials like silicon and glass (41), however, the new microwell chip consumables are more accessible, easy-to-use and disposable, making them suitable for a wider range of cell assay applications. In contrast to multi-layered microfluidic assemblies, our microwell chip is produced as a single unit that inherently has no risk of liquid leakage, issues with trapped air bubbles or requirement of external pumps and tubing. Long-term cell cultures are readily achieved due to the deep well geometry, since the culture medium can be exchanged by manual pipetting without disturbing the cells inside the wells. This is especially advantageous for long-term culturing of suspension cells, and we further leveraged on this potential by creating clonal expansions from single suspension or adherent cells. Furthermore, we established a straightforward protocol to create hundreds of uniformly sized spheroids in the microwells, with a minimal number of pipetting steps. The flat-bottom wells and COC plastic material were selected to offer an optical quality comparable to microscopy-grade glass coverslips and with maintained imaging access even during continued 2D or 3D cell culture. The bottom area of the wells is small enough to fit within a typical microscope field-of-view, facilitating extended single-cell tracking studies of even highly motile cells. All these properties make the chip an ideal platform for microscopy-based assays, especially when working with small cell numbers as is sometimes desired when working with patient material.

We also demonstrate the possibility to design complex workflows where initial time lapse imaging is combined with subsequent high-or even super-resolution imaging. Thus, this opens for new experimental approaches where dynamic parameters, such as serial killing at the single-cell level, or 3D spheroid killing at the population level, are related to for example lytic granule distribution and content in individual NK cells.

In addition to what has been demonstrated here, other types of analyses and read-outs (on-or off-chip) can also be applied and benefit from what the plastic microwell chips provide. Examples include genetic, proteomic and biochemical analyses that can be performed on cells that have grown and been treated in the chip. One specific example is flow cytometry that allows analysis of phenotype and viability of tumor cells from spheroids generated and treated in the chip, or immune cells that have infiltrated the spheroids. By using 16-chamber chips with the 8-slot microplate holder, it would for example be possible to test spheroids from 128 different conditions, with 36 spheroid replicates in each condition and a total number of 4 608 spheroids in the experiment. These analyses could also be performed in a complex workflow where initial imaging is followed by subsequent analysis.

Here, we demonstrated the potential of our method and microwell consumable for single-cell studies of immune cell function, by examining NK cell anti-tumor responses after treatments mimicking the tumor microenvironment (low glucose level, glutamine restriction, high lactate level and low oxygen level). Across our experiments, glutamine restriction appeared to have the strongest detrimental effect on NK cell responses, consistent with the proposed role of glutamine in sustaining cMyc activity and thereby effector functions in activated NK cells (51). A recent study showed that NK serial killing can increase upon long-term (3 weeks) treatment with the glucose transport inhibitor Glupin, likely due to a metabolic switch (50). Here we observed a slight decrease in NK serial killing after the 20 hour-treatment in low glucose. This difference could be due to a slow adjustment of the NK metabolism to low glucose (50), possibly related to expansion of specific NK cell subsets during long term culture. Interestingly, when cells were treated with both Glupin and glutaminase inhibitors the NK cells proliferated poorly and started to die within 6 days (50). This highlights the importance of studying the relative effects of various TME factors in conjunction, a research area in which there is currently a shortage of studies (52). To explore the relative effects of various TME factors on NK cell function, we used the multiplexing capacity of the 16-chamber to combine four different factors. Several combinations had an additive effect, however we found to our surprise a non-significant trend that some restrictions could benefit from being further combined with a high lactate concentration. This observation suggests that NK cells could potentially use lactate as an alternative energy source under restrictive conditions, as has previously been demonstrated for regulatory T cells (53), providing further support for the existence of redundancies in NK cell metabolism.

In summary, the presented plastic microwell chips support a range of complex imaging-based assays using both 2D and 3D cell cultures. The format and optical properties of the chips allow imaging across scales and magnifications. Building on that, we introduced a methodology for moving between rapid low-resolution viability screening, single-cell tracking and high-resolution assessment of intracellular protein expression in continuous experimental workflows, enabling correlative imaging of functional responses and phenotype. The microwell chips demonstrate the translation of specialized research technology to an easy-to-use but high-performance consumable for immunology applications.

## Limitations of the study

Injection molding requires careful design and specialized equipment for production of high-precision thermoplastic parts, which limits the flexibility of the technique. It is not a suitable approach for experimental production of small numbers of devices. The method does not readily allow the production of features below a few hundred micrometers, and smaller dimensions increase the risk of production defects, such as incomplete mold filling or weld lines during molding. The particular chips and protocols used here resulted in some undesired properties. The chips contain some weld lines, but they are mainly seen between microwells and they are only superficial so the effect is more cosmetic than functional. A general problem with relatively deep microwells is that the walls can give rise to shadowing effects causing uneven transmitted light images. However, this effect is even bigger when using non-transparent materials such as silicon. The plastic material and microwell array format are not compatible with some advanced imaging techniques, e.g. lattice light sheet microscopy. Chips that have been surface-treated with oxygen plasma become hydrophilic, which increases the risk for cross-contamination between chambers due to strong wetting behavior of aqueous liquids. On the contrary, chips that have been surface treated with non-adherence coating become hydrophobic, which increases the risk of unsuccessful wetting and filling of microwells with aqueous liquids. A small number of adherent cell lines that we tested showed a tendency to migrate also on the microwell walls affecting the imaging quality and quantification accuracy. The relatively small well size used here limits the sizes of spheroids or organoids that can be studied.

## Author contributions

H.v.O. and Q.V. designed the study, developed protocols, performed experiments, analyzed data and wrote the article. H.Z. developed image analysis pipelines, analyzed data and helped writing the article. P.S. developed vectors, transfected tumor cell lines and performed experiments. V.C. and K.O. developed protocols. T.F. performed experiments. A.K.W. developed vectors. N.S. designed the study, performed experiments, supervised the work and wrote the article. B.Ö. designed the study, supervised the work and wrote the article.

## Acknowledgements

We thank the group of Andreas Lundqvist at the Department of Oncology-Pathology, Karolinska Institutet for the for the cell lines ACHN and A498, the group of Per Kogner at the Childhood Cancer Research Unit at the Department of Women’s and Children’s Health, Karolinska Institutet for the cell line IMR-32, the group of Janne Lehtiö at the Department of Oncology and Pathology, Science for Life Laboratory, Karolinska Institutet for the cell line Sup-B15, the group of Anna Lundberg at Science for Life Laboratory, KTH Royal Institute of Technology for the cell line Jurkat, the group of Mikael Uhlin at the Department of Clinical Immunology and Transfusion Medicine at the Karolinska University Hospital for the cell line Kasumi-1, the group of Karl-Johan Malmberg at the Center for Infectious Medicine, Karolinska Institutet, and the Department of Cancer Immunology, Oslo University Hospital, for the cell line A549.

The LeGO-G2-puro was a gift from Boris Fehse at the Department of Stem Cell Transplantation, University Medical Center Hamburg-Eppendorf. The pLenti-CMV-MCS-RFP-SV-puro was a gift from Jonathan Garlick and Behzad Gerami-Naini. The pLJM1-EGFP was a gift from David Sabatini. The pJML1-NES-LQDT-mGFP-P2A-NES-VGPD mCherry and pLenti-CMV-MCS-RFP-SV-puro viral particles were produced by the VirusTech core facility at the Karolinska Institute.

We thank Rolf Nybom at SEMAIR Diagnostics AB, Sweden, for helping with Scanning Electron Microscopy; we acknowledge Hans Blom at the Advanced Light Microscopy unit at Science for Life Laboratory and the National Microscopy Infrastructure, NMI (VR-RFI 2016-00968) for providing assistance with STED imaging; we thank Brennan Wadsworth and Randall S. Johnson at the Department of Cell and Molecular Biology, Karolinska Institutet, for helping us perform hypoxia pretreatments.

We thank The Knut and Alice Wallenberg Foundation (Grant No 2018.0106), The Swedish Research Council (Grant No 2019-04925), The Swedish Foundation for Strategic Research (Grant No SBE13-0092), The Swedish Childhood Cancer Foundation (Grant No MT2019-0022, MT2022-0011), The Swedish Cancer Foundation (Grant No 19 0540 Pj) and Vinnova (Grant No 2019-00056) for financial support.

The schematic illustrations in Figures 1, 4 and 5 were created using Biorender.

## Materials and Methods

### Chip design and fabrication

The thermoplastic microwell chips were designed by us and manufactured by Tingverken AB, an industrial collaborator. The chips were produced using injection molding with Zeonor 1420R (COP, n=1.53), Topas 5013-L10 (COC, n=1.533) or Styrolution 124N/L (PS, n=1.56). The manufacturing first included flow simulations of the mold filling of the thermoplastic materials using the simulation software Moldex3D. Thereafter, the molds were produced by high-speed milling, wire-EDM and EDM. The 4-chamber version of the chip was produced as a first prototype using an Aluminum mold with lower manufacturing standards, whereas the 16-chamber version was produced using a hardened steel mold with higher surface finish and quality. The difference between the two versions can for example be observed in microscopy images of single wells (Fig. 1 G-H), where the 16-chamber version features more circular well shapes and smoother well surfaces and edges. Nonetheless, both versions are performing well for live cell assays.

### Liquid exchange and bead retrieval

TetraSpeck beads (Invitrogen) with a diameter of 4 μm were seeded in an untreated 16-chamber microwell chip containing 80 ul of PBS supplemented with 2% FBS and 2 mM EDTA (MACS buffer). The MACS buffer was removed and added 10 times using tilted pipetting, and after each exchange, an image of the chip was acquired using a 10x objective (10x/0.5 Plan-Apochromat) mounted on an AxioObserver7 wide-field microscope equipped with Colibri 7 LED illumination (Carl Zeiss AG). At the end, all beads were retrieved by flushing out the wells using vertical pipetting.

### Optical characterization

Resolution measurements were performed using TetraSpeck beads (Invitrogen) which exhibit 4 separate fluorescent emission peaks between 370 – 650 nm. The beads were seeded in the plastic chip chamber, on the surface of a #1.5 glass coverslip and in 10 wells distributed at different regions of a 96 well plate (Corning). According to the manufacturer’s instructions, the bead solution was then left to dry at room temperature in the dark before imaging. 3-color stack images of the beads were acquired using either a long-working distance 40x (LD Achroplan 40x/0.60, Carl Zeiss AG) or an oil-immersion 63x objective (Plan-Apochromat 63x/1.40, Carl Zeiss AG), mounted on a LSM 880 confocal microscope (Carl Zeiss AG). The PSF were then smoothed with a 2×2 median filter and fitted to Gaussian intensity profiles using a MATLAB script (54). Chromatic aberration was estimated in the datasets acquired with a long-working distance 40x objective. The focus plane of each color channel was found by performing a Gaussian fit on the variance-transformed fluorescent image and calculating the axial distance between respective focus planes. For transmission measurements, 4-color wide-field stack images of 4 µm-diameter TetraSpeck beads were acquired with a 10x objective (10x/0.5 Plan-Apochromat) mounted on an AxioObserver7 wide-field microscope equipped with Colibri 7 LED illumination (Carl Zeiss AG). The intensity for each bead was measured as the cumulated fluorescent intensity collected over all z-planes, and normalized to the mean intensity measured for beads imaged through a glass coverslip.

### Cell culture, vectors and transduction

#### Lentivirus production and titration

Lentivirus was produced as previously described (55). Briefly, 14×10^6^ HEK293FT cells were plated into a poly-D-lysine-coated 150-mm dish (Corning). The following day, the cells were transfected with 30 µg of LeGO-G2-puro (http://www.lentigo-vectors.de/vectors.htm), 15 µg of pMDLg/pRRE (#12251 Addgene, Cambridge, MA), 10 µg of pRSV-REV (#12553 Addgene), and 5 µg of phCMV-VSV-G (#8454 Addgene) using calcium phosphate transfection kit (Sigma-Aldrich) in the presence of 25 µM chloroquine (Sigma-Aldrich). The medium was changed 16 hours post-transfection, and the virus particles were collected after an additional 24 hr, by filtering the supernatant through a 0.45 µm filter. Virus particles were concentrated using lenti-X concentrator (Takara), and stored at -80°C. The virus was titrated by infection of HEK cells with subsequent detection of GFP+ cells by flow cytometry.

#### Tumor cell line transduction

K562 wild type cells were transduced at multiplicity of infection (MOI) 5 for 6 hours (K562-GFP) with 8 µg/ml protamine sulfate (Sigma Aldrich). After transduction, the cells were selected using 2 µg/ml puromycin (Thermo Fisher Scientific) and GFP expression was confirmed using flow cytometry. DLD-1 wild type cells were transduced with pLenti-CMV-MCS-RFP-SV-puro (Addgene plasmid #109377) viral particles (DLD-RFP) as described elsewhere (56). Both cell lines were transduced at low passage at MOI 3. The transduced cells were selected using 2 μg/mL of puromycin (Thermo Fisher Scientific).

#### Tumor cell line culture

Kidney cancer A498, Ovarian cancer OVCAR-8, Colon cancer DLD-1 (ATCC), kidney cancer ACHN, chronic myelogenous leukemia K562 (ATCC), acute T cell leukemia Jurkat, acute lymphocytic leukemia Nalm-6 (DSMZ), acute lymphocytic leukemia SUP-B15 and acute myeloblastic leukemia Kasumi-1 were grown in L-glutamine containing RPMI (Gibco) supplemented with 10% heat-inactivated fetal bovine serum (Sigma Aldrich) and 1% Penicillin/Streptomycin (Sigma Aldrich) (cRPMI). Breast cancer MCF-7 (ATCC) was grown in L-glutamine containing RPMI supplemented (Gibco) with 10% heat-inactivated fetal bovine serum (Sigma Aldrich), 1% Sodium Pyruvate (Sigma Aldrich) and 1% Penicillin/Streptomycin (Sigma Aldrich). Cervical cancer Hela (ATCC) was grown in L-glutamine containing DMEM (Gibco) supplemented with 10% heat-inactivated fetal bovine serum (Sigma Aldrich) and 1% Penicillin/Streptomycin (Sigma Aldrich). Neuroblastoma IMR-32 was grown in L-glutamine containing IMDM (Gibco) supplemented with 10% heat-inactivated fetal bovine serum (Sigma Aldrich) and 1% Penicillin/Streptomycin (Sigma Aldrich) (cIMDM). A549.Histone.mKate2 (A549), DLD-1-RFP and K562-GFP were cultured in cRPMI supplemented with 1 µg/ml Puromycin (Thermofisher). Adherent cell lines were split every 2-4 days using Trypsin/EDTA (Sigma Aldrich). Media was renewed for all cultures and experiments every 48 hours.

#### Primary cell culture

Human peripheral blood mononuclear cells (PBMCs) and NK cells were isolated from blood from anonymous healthy donors according to local ethics regulations. PBMCs were separated from buffy coats by density gradient centrifugation (Ficoll Paque, GE Healthcare) and stored at -140°C. NK cells were directly isolated from blood using negative selection (STEMCELL, EasySepTM, Direct Human NK Cell Isolation Kit). For TME treatment experiments, NK cells were cultured in U-bottom 96-well plates in cRPMI containing 100 U/ml of IL-2 (Peprotech) for 2 days, after which they were washed once in PBS and once in DMEM (no glucose, no glutamine, no phenol red Gibco) supplemented with 10% heat-inactivated, dialyzed FBS (Gibco), 1% Penicillin/Streptomycin (Sigma Aldrich) and 100 U/ml IL-2 (Peprotech) (Restricted Media). NK cells were resuspended in Restricted Media supplemented with either: for low glucose levels; 0.3 mM D-Glucose (Sigma Aldrich), 4.7 mM D-Mannitol (Sigma Aldrich), 2 mM L-glutamine (Sigma Aldrich), for glutamine restriction; 5 mM glucose (Sigma Aldrich), for high lactate; 5 mM glucose (Sigma Aldrich), 2 mM L-glutamine (Sigma Aldrich), 10 mM Lactic acid (Sigma aldrich), and for control; 5 mM glucose (Sigma Aldrich), 2 mM L-glutamine (Sigma Aldrich). For hypoxia treatment, the cells were placed in a hypoxia chamber with an oxygen level of 1%, or a normal incubator with an oxygen level of ∼20%. NK cells were treated in TME treatments for 24 hours.

### Chip preparation

#### Sterilization

Prior to any experiments, the chips were sterilized using 70% ethanol for 20 min. Ethanol-soaked chips were placed in a vacuum pump for the first few minutes of sterilization to ensure complete wetting of the wells. After sterilization, chips were left to dry in a sterile culture hood. Large batches of dry, sterile chips were stored under sterile conditions for future use. On the day of an experiment, surface treatment of the chips was performed.

#### Low-pressure oxygen plasma treatment

For 2D cell culture, improved cell binding to the surface was obtained by exposing the chips to an oxygen plasma at a pressure of 7×10^−2^ bar for 30 s (CUTE, Femto Science Inc). For this study, plasma-treated chips were used within 12 hours, but chips maintained the cell-adhesive properties for at least 20 days. Before cell seeding, complete medium was added to the dry and sterile plasma-treated or untreated chips, which were then centrifuged at 1200 g for 30 s to ensure wetting of all microwells.

#### Anti-adherence treatment

When preparing chips for the generation of 3D spheroids, the surface of untreated chips was instead coated with a protein-repellent anti-adherence rinsing solution (STEMCELL Technologies). The solution was directly added to dry and sterile chips, followed by centrifugation at 1200 g for 30 s to ensure wetting of the microwells. The rinsing solution was left to incubate at room temperature for 15 min after which it was removed, and the chips were thoroughly washed three times with warm complete medium before cells were added.

### On-chip 2D and 3D cell culture

#### Cell seeding and cell counting

Tumor cells were harvested from their culture vessel, and washed once in complete medium. Depending on the division rate of each cell type, 10 000-30 000 cells were added to one chamber of a 4-chambered chip (resulting in approximately 20-60 cells per well). For clonal experiments, 1000 cells were instead added. Cells were left to seed into the wells for 30 min, after which the chip was washed 3 times to ensure that cells that were located in between the wells were removed. Pipetting was done into the corners of the chambers with an angled pipetting technique (∼45 degrees) to avoid disturbing the cells within the wells. The chip was covered with a lid and placed in the incubator. For 3D culture, the chip was either placed in a horizontal position, or angled at approximately 15 degrees with the help of a wedge. After 24 hours of culture, the chip was again placed in a horizontal position. Medium was completely exchanged on all chips every 48 hours by pipetting at an angle into the corner.

For all cell division studies, a starting image was acquired of the cells right after seeding (and before tilting of the chip in the case of 3D culture). This enabled the counting of cell numbers in each well at seeding. Wide-field imaging was performed using a Zeiss Axio Observer 7 microscope (Carl Zeiss AG) equipped with a 10x/0.5 Plan-Neofluar objective and a Hamamatsu ORCA-flash 4.0 LT camera, or one equipped with a 10x/0.5 Plan-Apochromat objective and a Hamamatsu ORCA-flash 4.0 camera (both microscopes are referred to as a wide-field microscope). Both microscopes had an incubation chamber with environmental control (37°C, 5% CO_2_).

#### 2D cell culture of cancer cell line and primary immune cells

The division rate of five adherent and five suspension cell lines was measured by culturing cells in plasma-treated chips in an incubator for 48 hours. After this, all culture medium was removed and PBS containing 5 μM DRAQ5 (BD Biosciences) and 4 μM Abberior LIVE 590 tubulin (Abberior) was added to the chip. The chip was incubated at 37°C for 30 min and images were then acquired on a confocal Zeiss LSM 880 equipped with a 20x/0.8 Plan-Apochromat objective and an incubation chamber. The nuclear stain was used to manually calculate the number of cells in each well at 48 hours, and cell division, determined as the fold change from the number of cells at seeding, was determined as the average of 2-5 microwells per cell line.

The division kinetics of DLD-1-RFP and K562-GFP cells were measured by culturing the cells in a plasma-treated chip on the wide-field microscope for 72 hours, and acquiring images every 30-60 min. Simultaneous imaging of cells seeded at a similar density in a cell culture-treated 96-well plate (Corning) was performed for comparison. The cells were automatically segmented based on their fluorescent signal using the pixel classification module in the software ilastik (57). Briefly, a small subset of the data was manually annotated to train the neural network until satisfying segmentation accuracy was reached, after which the pipeline was applied to the rest of the data. A subset of the data was visually inspected to confirm the adequacy of the segmentation. The confluency of cells in a well was determined as the total area of cells in that well, divided by the average area of a well. For cells grown on the 96-well plate, confluency was determined as the total area of cells in one image divided by the area of the image. For the analysis of spheroid formation dynamics, the spheroids were grown directly on the wide-field microscope (horizontally positioned), with and without anti-adherence coating. Images of the chip were acquired every 60 minutes, and the spheroids were automatically detected using the pixel classification module in ilastik, followed by area measurements in ImageJ. The average number of spheroids and the average spheroid area was calculated for each well.

For clonal expansions and primary cell culture experiments, the cells were grown in plasma-treated chips for 7-14 days while placed in an incubator. Every 2-3 days, the chip was moved to a wide-field microscope and images of the chip were acquired. Clonal expansion rates of 2D cultured DLD-1 and K562 cells were determined based on their confluency as described above. Before seeding on the chip, NK cells were stained with 5 μM CellTrace Yellow (Thermofisher) in PBS for 10 min. The stained NK cells, or unstained PBMCs were seeded at low density on a plasma-treated chip containing cIMDM supplemented with 10 ng/ml IL-15 (PeproTech) and 100 U/ml IL-2 (PeproTech). NK cell division was determined as the number of cells in a well divided by the number of cells in the well after seeding.

#### 3D cell culture of cancer cell lines

For the investigation of cell division of DLD-1 cells grown in 3D, spheroids were formed and cultured for 48 hours. After this, the spheroids were washed three times with PBS, followed by a 15 min fixation with Cytofix/Cytoperm (BD Biosciences). Fixed spheroids were washed three times with washing buffer (PBS supplemented with 0.2% Triton X-100, 0.05% Tween-20 and 2% w/v bovine serum albumin, BSA), then left in staining buffer (PBS, 0.2% Triton X-100, 0.05% Tween-20 and 0.1% w/v BSA) for 15 min. The spheroids were incubated with blocking buffer (PBS, 0.2% Triton X-100, 0.05% Tween-20, 0.1% w/v BSA and 5% goat serum) for 1 hour, followed by an overnight incubation at 4°C with staining buffer (nuclear staining) or a 36-hour incubation at room temperature with staining buffer containing 2.5 µg/mL of an antibody against vimentin (ab195877, abcam; nuclear and extracellular staining). The spheroids were then washed 3 times with the washing buffer and incubated with PBS containing 20 µM DRAQ5 for 4 hours, followed by 3 steps of washing with the washing buffer. The stained spheroids were mounted in a refractive index-matching medium, by gradually increasing the concentration (0, 10, 30, 50, 80, 100%) of Histodenz in PBS. 3D confocal stacks were acquired on a Zeiss LSM 880 equipped with a 20x/0.8 Plan-Apochromat objective (Carl Zeiss AG) using the FAST AiryScan module with a z-step of 0.5 µm. The number of cells within each spheroid at 48 hours was determined by automatic detection of nuclei using Imaris. The total volume of the spheroid was estimated by automatic detection of a surface surrounding all nuclei of the spheroid. Cell division was calculated as a fold change from the number of cells manually counted in the image taken after seeding.

Clonal spheroids were formed from single DLD-1-RFP cells by seeding 1000 cells into a chamber of an anti-adherence-coated chip. The chip was imaged every 7 days and following 14 days of cell culture in the incubator, the size of clonally formed spheroids was determined by manual measurement of the spheroid diameter and an average number was calculated for each well.

### Super-resolution imaging

Plasma-treated plastic chips were coated with antibodies against NK cell activating receptors LFA-1 (BioLegend, clone HI111) and CD16 (BioLegend, clone 3G8) diluted at 2 µg/mL in PBS for 4 hours. After washing the chips, NK cells were seeded into the wells and left to interact with the well surface for 30 min. The cells were then fixed and permeabilized using Cytofix (BD Biosciences) for 15 min, washed 3 times using Perm/Wash (BD Biosciences) before blocking with PBS, 5% goat serum and 0.5% w/v BSA. The cells were then labeled for F-actin using phalloidin-STAR Red (abberior) at 2 µg/mL, and for PKC-Θ using a primary rabbit antibody at 2 µg/mL (C-18, polyclonal, Santa Cruz Biotechnology) overnight at 4°C. Then, a secondary staining was performed, using an Alexa Fluor 488-conjugated goat anti-rabbit antibody at 4 µg/mL (ab150089, abcam) for 1 hour at room temperature. The samples were mounted in ProLong Gold (Thermo Fisher Scientific) and kept in the dark at 4°C until imaging. Confocal and STED images were acquired with a 100x/1.4 oil-immersion objective (Leica Microsystems) installed on a Leica SP8 3X STED microscope. Stimulated depletion was induced using a laser at 592 nm wavelength.

### Immune cell killing in 2D and 3D

#### 2D experimental layout

For 2D killing assays, A549.Histone.mKate2 (A549) cells were seeded in a plasma-treated chip containing cRPMI culture medium, then left to adhere for 18 hours. NK cells were harvested and pretreated in four conditions reproducing the TME (see “Primary cell culture”). After 24 hours of treatment, the NK cells were washed once in PBS and thereafter stained with 5 μM CellTrace Yellow in PBS for 10 min at 37°C. Phenol-free cRPMI containing 100 U/ml of IL-2 (Peprotech), 1 µg/ml of Cetuximab (Invivogen) and 200 ng/mL Sytox Green was added to the chip containing the tumor cells which were left to incubate for 30 min. After this, stained and washed NK cells were added to the chip resulting in an average of 6 NK cells per well. The chip was thoroughly washed 3 times to remove NK cells lying between the wells. The chip was then placed on a wide-field microscope and images were acquired every 3 min for 5 hours. After 5 hours of co-cultures, the chip was taken off the microscope and immediately washed 3 times with PBS, before fixation with Cytofix (BD Biosciences) at room temperature for 15 min. The cells were washed and permeabilized 3 times 5 min in Perm/Wash (BD Biosciences), followed by a 1 hour blocking step at room temperature with PBS + 5% FBS + 0.5% BSA. The cells were washed again with Perm/Wash for 3 times 5 minutes, then stained with an antibody against Granzyme B (clone GB11), directly conjugated with Alexa Fluor 647 (BD Biosciences) at 2.5 µg/mL in Perm/Wash, for 18 hours at 4°C. After 3 times 5 minutes washes with Perm/Wash followed by 3 times 5 minutes in PBS, the co-cultures were mounted in ProlongGold and kept to cure in the dark at 4°C until imaging.

#### 3D experimental layout

For 3D killing assays, A549 spheroids were created in the chips 72 hours before the day of the experiment. 24 hours before the experiment, NK cells were harvested and pretreated in 16 conditions reproducing the TME (see “Primary cell culture”). On the day of the experiment, spheroids were incubated with cRPMI containing 1 µg/mL of Cetuximab (Invivogen) and 500 ng/mL Sytox Green for 30 minutes. Meanwhile, NK cells were stained as described in the *2D experimental layout*. Stained NK cells were seeded to the chip resulting in ∼20-60 NK cells per well. The chip was thoroughly washed 3 times to remove NK cells that were lying between the wells. The chip was then placed on a wide-field microscope and images were acquired every 15 minutes for 16-20 hours. After 16 hours, the spheroids were thoroughly washed three times with PBS, followed by a 20 minutes fixation using Cytofix (BD Biosciences). Then, the spheroids were incubated 15 minutes in the staining buffer described above, followed by 3 hours in blocking buffer. The spheroids were labeled with an antibody against Granzyme B (clone GB11), directly conjugated with Alexa Fluor 647 (BD Biosciences) at 2.5 µg/mL in the staining buffer, for 36 hours at room temperature. Unbound antibodies were washed off for 3 times 20 minutes in the washing buffer, then the co-cultures were mounted in refractive index-matching medium, by the addition of a stepwise increasing concentration (0, 10, 30, 50, 80, 100%) of Histodenz in PBS. 3D confocal images were acquired on the confocal system using the FAST AiryScan module with a z-step of 2 µm.

#### Image analysis

For analyzing killing assays, we trained state-of-the-art convolutional neural networks based on resnet50 as a backbone structure to conduct automated image segmentation and analysis (58). We used two deep neural networks to find an accurate number of dead tumor cells (2D only) and live NK cells (both 2D and 3D) in each well, based on the fluorescence and bright-field information (Supp. Movie 2). We observed that tumor cells did not always take up the fluorescent death marker in the 5 hours assay, yet they showed clear morphological changes (membrane blebbing, shrinkage or swelling) as a result of cell death that could be recognized in the bright-field channel. Therefore, we used combined fluorescence and bright-field information to segment live and dead cells.

To implement the networks adapted to our data, we designed the network input as 261 by 261 images with 4 channels, including 3 fluorescence channels and 1 bright-field channel. First, we trained an NK cell segmentation network to output four classes, corresponding to living cells, dead cells, cell contour and background pixels. The training data contained images of NK interacting with both spheroid and individual tumor cells. A separate network was trained to focus on finding dead tumor cells in 2D cytotoxicity data. We annotated 500+ samples for the training of both networks. The two networks achieved 99.4% and 99.6% classification accuracy for our NK cells and dead tumor cells, respectively. Further, we evaluated the accuracy of the network using the number of detected cells compared to the ground truth and the results showed 86.2% and 89.7% accuracy using a local test dataset.

For 2D killing assays, a killing index was obtained by relating the dead tumor cell count according to our neural network, to the NK cell count in the well at the start of the assay, averaged in the first two frames of the time-lapse dataset to account for segmentation errors. The serial-killing behavior of single NK cells was manually analyzed in each well. For 3D killing assays, tumor cell death in the spheroid was estimated by measuring the cumulated Sytox fluorescent intensity in each well, removing the intensity at the start of the assay. For correlation between infiltration and killing, the exact number of dead cells was detected by segmenting sytox positive objects in the 3D images using Imaris. Dead NK cells were excluded. For all analyses involving the comparison of TME treatments, the wells were selected so that the mean number of live NK cells per well was equal between the conditions. The target cells were in large excess in both 2D and 3D experiments.

#### Resource availability

The aim is to make the microwell chips available for external users, either as test samples or for a fee. Please, contact the lead author for further information.

## References

1. Chaplin DD. Overview of the immune response. J Allergy Clin Immunol. 2010 Feb;125(2 SUPPL. 2).

2. Meuer SC, Fitzgerald KA, Hussey RE, Hodgdon JC, Schlossman SF, Reinherz EL. Clonotypic structures involved in antigen-specific human T cell function: Relationship to the T3 molecular complex. J Exp Med. 1983 Feb 1;157(2):705–19.

3. Hibi T, Dosch H -M. Limiting dilution analysis of the B cell compartment in human bone marrow. Eur J Immunol. 1986;16(2):139–45.

4. Sedgwick JD, Holt PG. A solid-phase immunoenzymatic technique for the enumeration of specific antibody-secreting cells. J Immunol Methods. 1983 Feb 25;57(1–3):301–9.

5. Deniz G, Erten G, Kücüksezer UC, Kocacik D, Karagiannidis C, Aktas E, et al. Regulatory NK Cells Suppress Antigen-Specific T Cell Responses. J Immunol. 2008 Jan 15;180(2):850–7.

6. Paul S, Kulkarni N, Shilpi N, Lal G. Intratumoral natural killer cells show reduced effector and cytolytic properties and control the differentiation of effector Th1 cells. OncoImmunology. 2016 Dec 1;5(12).

7. Bengsch B, Ohtani T, Herati RS, Bovenschen N, Chang KM, Wherry EJ. Deep immune profiling by mass cytometry links human T and NK cell differentiation and cytotoxic molecule expression patterns. J Immunol Methods. 2018 Feb 1;453:3–10.

8. Ni J, Wang X, Stojanovic A, Zhang Q, Wincher M, Bühler L, et al. Single-Cell RNA Sequencing of Tumor-Infiltrating NK Cells Reveals that Inhibition of Transcription Factor HIF-1α Unleashes NK Cell Activity. Immunity. 2020 Jun 16;52(6):1075–1087.e8.

9. Szabo PA, Levitin HM, Miron M, Snyder ME, Senda T, Yuan J, et al. Single-cell transcriptomics of human T cells reveals tissue and activation signatures in health and disease. Nat Commun. 2019 Dec 1;10(1).

10. Chokkalingam V, Tel J, Wimmers F, Liu X, Semenov S, Thiele J, et al. Probing cellular heterogeneity in cytokine-secreting immune cells using droplet-based microfluidics. Lab Chip. 2013 Dec 21;13(24):4740–4.

11. Bandey IN, Adolacion JRT, Romain G, Paniagua MM, An X, Saeedi A, et al. Designed improvement to T-cell immunotherapy by multidimensional single cell profiling. J Immunother Cancer. 2021 Mar 15;9(3).

12. Merouane A, Rey-Villamizar N, Lu Y, Liadi I, Romain G, Lu J, et al. Automated profiling of individual cell-cell interactions from high-throughput time-lapse imaging microscopy in nanowell grids (TIMING). Bioinformatics. 2015;31(19):3189–97.

13. Guldevall K, Vanherberghen B, Frisk T, Hurtig J, Christakou AE, Manneberg O, et al. Imaging immune surveillance of individual natural killer cells confined in microwell arrays. PLoS ONE. 2010;

14. Subedi N, Verhagen LP, de Jonge P, Van Eyndhoven LC, van Turnhout MC, Koomen V, et al. Single-Cell Profiling Reveals Functional Heterogeneity and Serial Killing in Human Peripheral and Ex Vivo-Generated CD34+ Progenitor-Derived Natural Killer Cells. Adv Biol. 2023;7(4):2200207.

15. Antona S, Platzman I, Spatz JP. Droplet-Based Cytotoxicity Assay: Implementation of Time-Efficient Screening of Antitumor Activity of Natural Killer Cells. ACS Omega. 2020 Sep 29;5(38):24674–83.

16. Guldevall K, Brandt L, Forslund E, Olofsson K, Frisk TW, Olofsson PE, et al. Microchip screening platform for single cell assessment of NK cell cytotoxicity. Front Immunol. 2016;7(APR):1–7.

17. Vanherberghen B, Olofsson PE, Forslund E, Sternberg-Simon M, Khorshidi MA, Pacouret S, et al. Classification of human natural killer cells based on migration behavior and cytotoxic response. Blood. 2013;121(8):1326–34.

18. Prager I, Liesche C, van Ooijen H, Urlaub D, Verron Q, Sandström N, et al. NK cells switch from granzyme B to death receptor–mediated cytotoxicity during serial killing. J Exp Med. 2019 Sep 2;216(9):2113–27.

19. Vivier E. Functions of natural killer cells. Nat Immunol. 2008;9(5):503–10.

20. Ruggeri L, Capanni M, Urbani E, Perruccio K, Shlomchik WD, Tosti A, et al. Effectiveness of donor natural killer cell aloreactivity in mismatched hematopoietic transplants. Science. 2002 Mar 15;295(5562):2097–100.

21. Dall’Ozzo S, Tartas S, Paintaud G, Cartron G, Colombat P, Bardos P, et al. Rituximab-dependent cytotoxicity by natural killer cells: Influence of FCGR3A polymorphism on the concentration-effect relationship. Cancer Res. 2004;64(13):4664–9.

22. Guillerey C, Huntington ND, Smyth MJ. Targeting natural killer cells in cancer immunotherapy. Nat Immunol. 2016 Aug 19;17(9):1025–36.

23. Liu S, Galat V, Galat4 Y, Lee YKA, Wainwright D, Wu J. NK cell-based cancer immunotherapy: from basic biology to clinical development. J Hematol OncolJ Hematol Oncol. 2021 Dec 1;14(1).

24. Raghavan S, Mehta P, Xie Y, Lei YL, Mehta G. Ovarian cancer stem cells and macrophages reciprocally interact through the WNT pathway to promote pro-tumoral and malignant phenotypes in 3D engineered microenvironments. J Immunother Cancer. 2019 Jul 19;7(1).

25. Chakrabarti J, Holokai L, Syu LJ, Steele NG, Chang J, Wang J, et al. Hedgehog signaling induces PD-L1 expression and tumor cell proliferation in gastric cancer. Oncotarget. 2018 Dec 1;9(100):37439–57.

26. Ayuso JM, Truttschel R, Gong MM, Humayun M, Virumbrales-Munoz M, Vitek R, et al. Evaluating natural killer cell cytotoxicity against solid tumors using a microfluidic model. OncoImmunology. 2019 Mar 4;8(3).

27. Kaur A, Ecker BL, Douglass SM, Kugel CH, Webster MR, Almeida FV, et al. Remodeling of the collagen matrix in aging skin promotes melanoma metastasis and affects immune cell motility. Cancer Discov. 2019 Jan 1;9(1):64–81.

28. Courau T, Bonnereau J, Chicoteau J, Bottois H, Remark R, Assante Miranda L, et al. Cocultures of human colorectal tumor spheroids with immune cells reveal the therapeutic potential of MICA/B and NKG2A targeting for cancer treatment. J Immunother Cancer. 2019 Mar 14;7(1).

29. Sherman H, Gitschier HJ, Rossi AE. A novel three-dimensional immune oncology model for high-throughput testing of tumoricidal activity. Front Immunol. 2018 Apr 23;9(APR).

30. Muranen T, Selfors LM, Worster DT, Iwanicki MP, Song L, Morales FC, et al. Inhibition of PI3K/mTOR Leads to Adaptive Resistance in Matrix-Attached Cancer Cells. Cancer Cell. 2012 Feb 14;21(2):227–39.

31. Langan LM, Dodd NJF, Owen SF, Purcell WM, Jackson SK, Jha AN. Direct measurements of oxygen gradients in spheroid culture system using electron parametric resonance oximetry. PLoS ONE. 2016 Feb 1;11(2).

32. Zboralski D, Hoehlig K, Eulberg D, Frömming A, Vater A. Increasing tumor-infiltrating T cells through inhibition of CXCL12 with NOX-A12 synergizes with PD-1 blockade. Cancer Immunol Res. 2017 Nov 1;5(11):950–6.

33. Carannante V, Wiklund M, Önfelt B. In vitro models to study natural killer cell dynamics in the tumor microenvironment. Front Immunol. 2023;14:1135148.

34. Booij TH, Price LS, Danen EHJ. 3D Cell-Based Assays for Drug Screens: Challenges in Imaging, Image Analysis, and High-Content Analysis. SLAS Discov Adv Life Sci R D. 2019 Feb 28;24(6):615– 27.

35. Bresciani G, Hofland LJ, Dogan F, Giamas G, Gagliano T, Zatelli MC. Evaluation of Spheroid 3D Culture Methods to Study a Pancreatic Neuroendocrine Neoplasm Cell Line. Front Endocrinol. 2019 Oct 4;10.

36. Richardson DS, Lichtman JW. Clarifying Tissue Clearing. Cell. 2015 Jul 18;162(2):246–57.

37. Amaral RLF, Miranda M, Marcato PD, Swiech K. Comparative analysis of 3D bladder tumor spheroids obtained by forced floating and hanging drop methods for drug screening. Front Physiol. 2017 Aug 22;8(AUG).

38. Mehta G, Hsiao AY, Ingram M, Luker GD, Takayama S. Opportunities and challenges for use of tumor spheroids as models to test drug delivery and efficacy. J Controlled Release. 2012 Dec 10;164(2):192–204.

39. Olofsson K, Carannante V, Ohlin M, Frisk T, Kushiro K, Takai M, et al. Acoustic formation of multicellular tumor spheroids enabling on-chip functional and structural imaging. Lab Chip. 2018;

40. Christakou AE, Ohlin M, Onfelt B, Wiklund M. Ultrasonic three-dimensional on-chip cell culture for dynamic studies of tumor immune surveillance by natural killer cells. Lab Chip. 2015;15(15):3222–31.

41. Sandström N, Carannante V, Olofsson K, Sandoz PA, Moussaud-Lamodière EL, Seashore-Ludlow B, et al. Miniaturized and multiplexed high-content screening of drug and immune sensitivity in a multichambered microwell chip. Cell Rep Methods. 2022 Jul;2(7):100256.

42. Bernard M, Jubeli E, Bakar J, Tortolano L, Saunier J, Yagoubi N. Biocompatibility assessment of cyclic olefin copolymers: Impact of two additives on cytotoxicity, oxidative stress, inflammatory reactions, and hemocompatibility. J Biomed Mater Res - Part A. 2017 Dec 1;105(12):3333–49.

43. Nunes PS, Ohlsson PD, Ordeig O, Kutter JP. Cyclic olefin polymers: Emerging materials for lab-on-a-chip applications. Microfluid Nanofluidics. 2010 Aug;9(2–3):145–61.

44. Jeon JS, Chung S, Kamm RD, Charest JL. Hot embossing for fabrication of a microfluidic 3D cell culture platform. Biomed Microdevices. 2011 Apr;13(2):325–33.

45. Al-Azzam N, Alazzam A. Micropatterning of cells via adjusting surface wettability using plasma treatment and graphene oxide deposition. PLOS ONE. 2022 Jun 16;17(6):e0269914.

46. Khan GN, Kim EJ, Shin TS, Lee SH. Heterogeneous Cell Types in Single-cell-derived Clones of MCF7 and MDA-MB-231 Cells. Anticancer Res. 2017 May 7;37(5):2343–54.

47. Locke M, Heywood M, Fawell S, Mackenzie IC. Retention of Intrinsic Stem Cell Hierarchies in Carcinoma-Derived Cell Lines. Cancer Res. 2005 Oct 1;65(19):8944–50.

48. Ferrand A, Sandrin MS, Shulkes A, Baldwin GS. Expression of gastrin precursors by CD133-positive colorectal cancer cells is crucial for tumour growth. Biochim Biophys Acta. 2009 Mar;1793(3):477–88.

49. Haroun-Izquierdo A, Vincenti M, Netskar H, van Ooijen H, Zhang B, Bendzick L, et al. Adaptive single-KIR+NKG2C+ NK cells expanded from select superdonors show potent missing-self reactivity and efficiently control HLA-mismatched acute myeloid leukemia. J Immunother Cancer. 2022 Nov;10(11):e005577.

50. Picard LK, Niemann JA, Littwitz-Salomon E, Waldmann H, Watzl C. Restriction of Glycolysis Increases Serial Killing Capacity of Natural Killer Cells. Int J Mol Sci. 2024 Mar 2;25(5):2917.

51. Loftus RM, Assmann N, Kedia-Mehta N, O’Brien KL, Garcia A, Gillespie C, et al. Amino acid-dependent cMyc expression is essential for NK cell metabolic and functional responses in mice. Nat Commun. 2018 Jun 14;9(1):2341.

52. Poznanski SM, Ashkar AA. What Defines NK Cell Functional Fate: Phenotype or Metabolism? Front Immunol [Internet]. 2019 [cited 2023 May 27];10. Available from: https://www.frontiersin.org/articles/10.3389/fimmu.2019.01414

53. Watson MJ, Vignali PDA, Mullett SJ, Overacre-Delgoffe AE, Peralta RM, Grebinoski S, et al. Metabolic support of tumour-infiltrating regulatory T cells by lactic acid. Nature. 2021 Mar 25;591(7851):645–51.

54. Campbell R. measurePSF [Internet]. GitHub; 2021. Available from: https://github.com/raacampbell/measurePSF

55. Weber K, Thomaschewski M, Benten D, Fehse B. RGB marking with lentiviral vectors for multicolor clonal cell tracking. Nat Protoc. 2012 May;7(5):839–49.

56. Barde I, Salmon P, Trono D. Production and Titration of Lentiviral Vectors. Curr Protoc Neurosci. 2010 Oct;

57. Berg S, Kutra D, Kroeger T, Straehle CN, Kausler BX, Haubold C, et al. Ilastik: Interactive Machine Learning for (Bio)Image Analysis. Nat Methods. 2019;16(12):1226–32.

58. He K, Zhang X, Ren S, Sun J. Deep Residual Learning for Image Recognition. In: 2016 IEEE Conference on Computer Vision and Pattern Recognition (CVPR) [Internet]. Las Vegas, NV, USA: IEEE; 2016 [cited 2024 Sep 26]. p. 770–8. Available from: http://ieeexplore.ieee.org/document/7780459/

