## Supplemental figures and tables for "A thermoplastic chip for correlative assays combining screening and high-resolution imaging of immune cell responses"

**Supporting figure S1.** A microwell array optimized for wide-field microscopy.

**Supporting figure S2.** Effect of the bottom thickness on the optical performance of the chip.

**Supporting figure S3.** 2D cell division in the microwell chip.

**Supporting figure S4.** Cell proliferation across the microwell chip

**Supporting figure S5**. Comparison of protocols to form spheroids in the microwells.

**Supporting figure S6.** Improvement to deep imaging of spheroids by optical clearing.

**Supporting Table 1**. Spheroid formation in microwells using a range of immortal cell lines.

**Supporting movie 1**. Successive liquid exchanges and bead retrieval in a microwell chip containing 4 µm-wide beads

**Supporting movie 2.** Example of killing events by an NK cell, illustrating how our machine learning algorithm could identify tumor cell death by uptake of fluorescent markers or by characteristic morphological changes.

**Supporting movie 3**. Correlative assay in the microwell chip, relating the cytotoxic potential of single NK cells against A549 targets, to their granzyme B content at the end of the assay.

**Supporting movie 4**. Correlative assay in the microwell chip, relating the infiltration of single NK cells into 3D spheroids of A549 tumor cells, to their granzyme B content at the end of the assay.

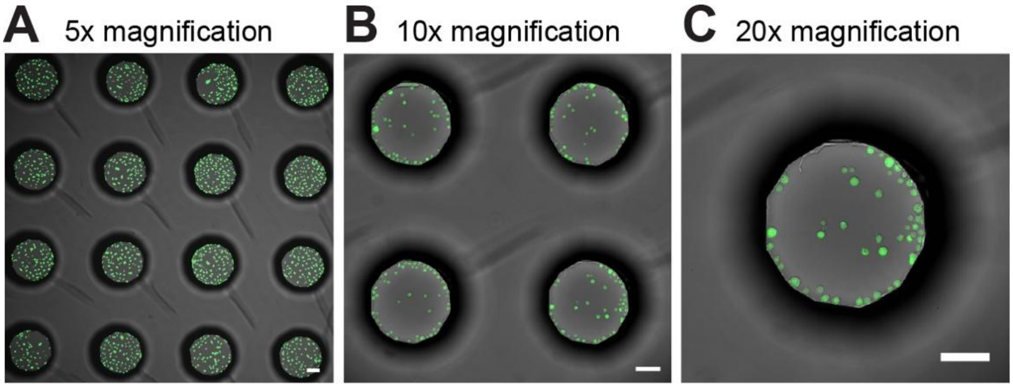


***Supporting figure S1. A microwell array optimized for wide-field microscopy.***
***(A-C)*** *Example wide-field microscopy images of the region of the chip fitting within the FoV using a 5x objective (A), a 10x objective (B) or a 20x objective (C), and a camera sensor of 13x13 mm$^2$. Scale bars: 100 µm.*

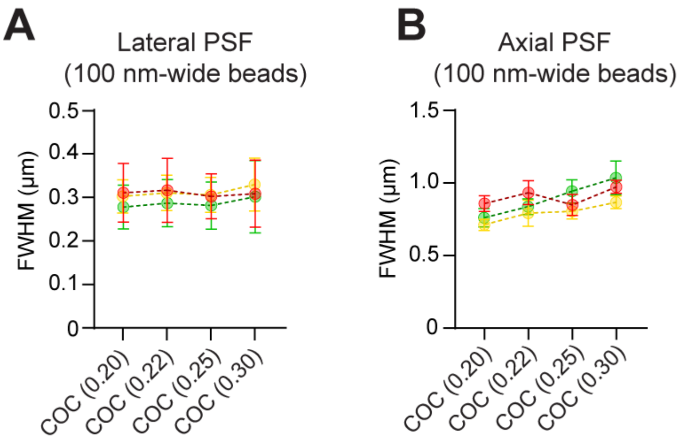


***Supporting figure S2. Effect of the bottom thickness on the optical performance of the chip.***
***(A-B)*** *Sub-diffraction 100 nm-wide beads seeded in the chip microwells were imaged with an oil-immersion 63x/1.4 objective. Width of the lateral (A) and axial (B) PSF. Pooled data from 3 independent experiments, with a total of n=94-129 beads per condition.*

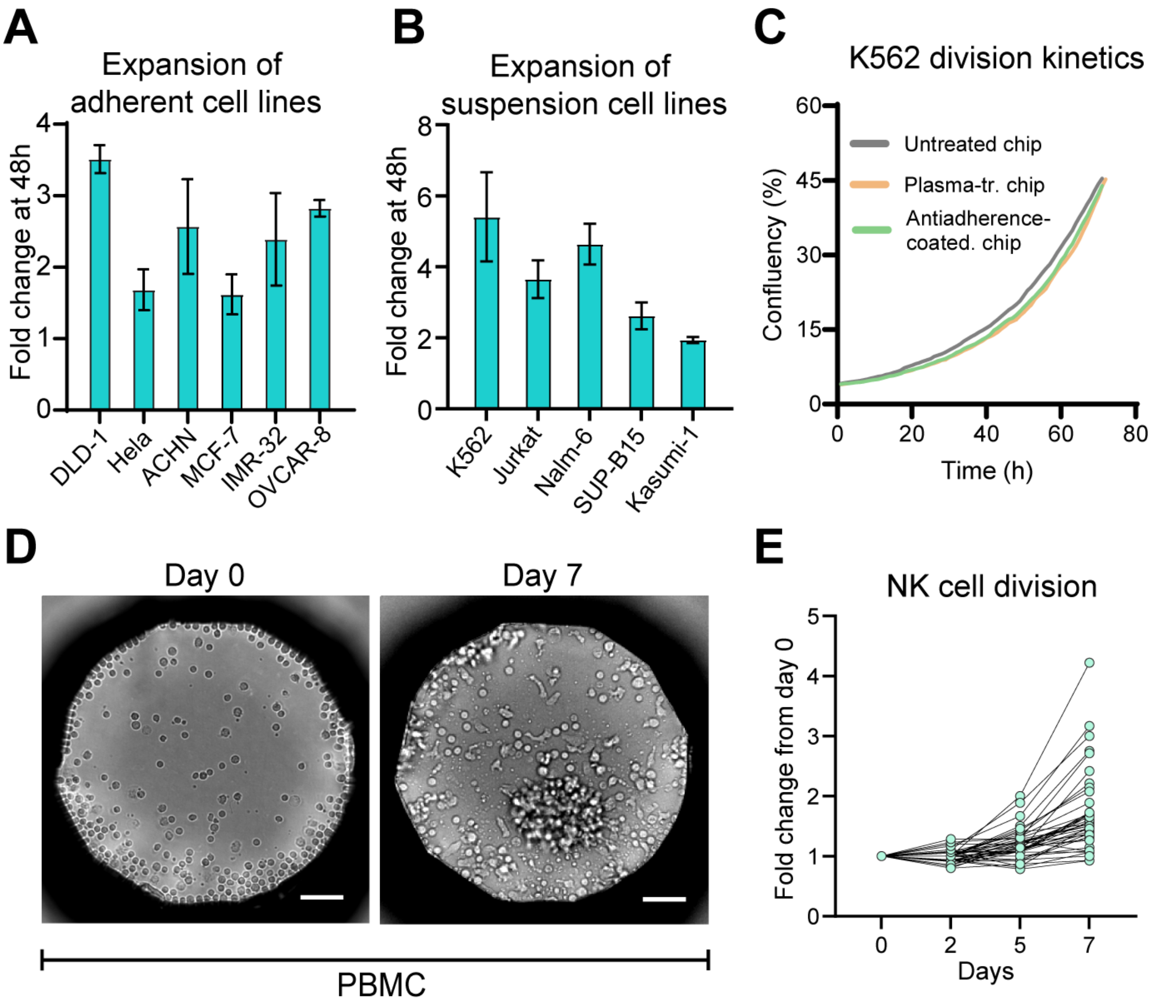


***Supporting figure S3. 2D cell division in the microwell chip.***

***(A)*** *Division rate of a range of adherent cell lines grown in oxygen plasma-treated microwells.* ***(B)*** *Division rate of a range of suspension cell lines grown in anti-adherence-treated microwells. Each bar in (A-B) represents the average of 2-5 wells with corresponding standard deviation.* ***(C)*** *Confluency over time, for K562 cells grown in microwells with different surface treatments.* ***(D)*** *Example images of PBMCs grown for 7 days in anti-adherence-treated microwells. Scale bars: 50 µm.* ***(E)*** *Division kinetics of primary NK cells grown for 7 days in anti-adherence-treated microwells.*

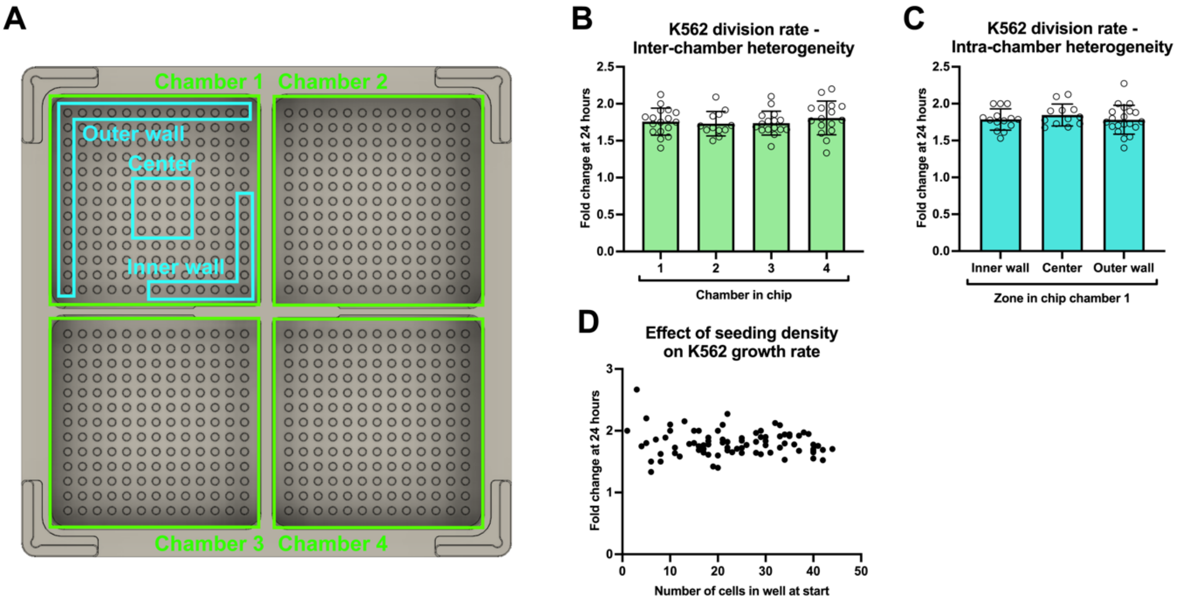


***Supporting figure S4. Cell proliferation across the microwell chip.*** *K562 tumor cells were cultured in the microwells, and their growth rate was related to the well location on the chip.* ***(A)*** *The microwell array, composed of 4 chambers, was further divided into regions of interest with possibly different environmental conditions.* ***(B-C)*** *Growth rate of K562 cells over 24 hours, compared between chambers 1-4 (B) or between regions of a single chamber (C).* ***(D)*** *Correlation between the average K562 growth rate in microwells and the number of cells at the start of the experiment. Each bar in (B-C) indicates the average of at least 8 microwells, while each dot in (B-D) represents a single microwell. Only wells containing at least 5 cells were included in the analysis in (B-C).*

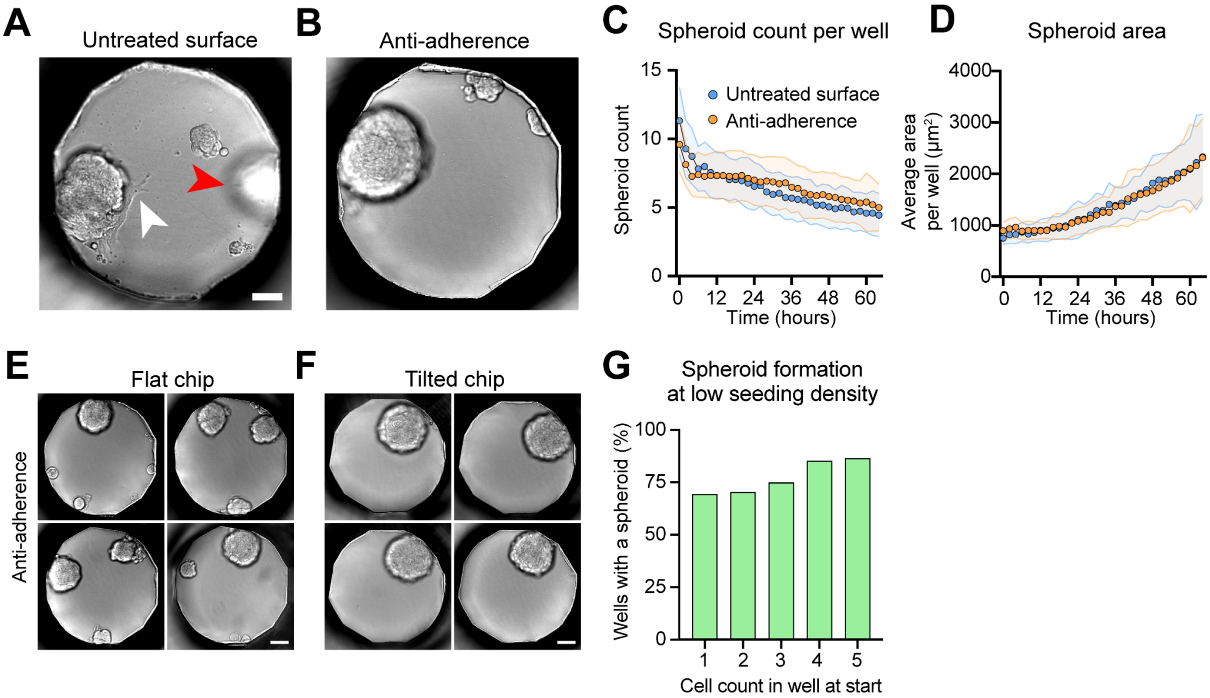


***Supporting figure S5. Comparison of protocols to form spheroids in the microwells.***
***(A-B)*** *Example bright-field images of DLD-1 spheroids formed in untreated (A) and anti-adherence-treated (B) microwells. White arrowhead: protrusions binding a spheroid to the untreated surface of the well. Red arrowhead: spheroid displaced to an out-of-focus plane.* ***(C-D)*** *Comparison of the growth kinetics of spheroids in untreated and anti-adherence-treated chips, considering the average number of cell clusters per well (C) and their mean area (D).* ***(E-F)*** *Representative bright-field images of the spheroids obtained by either keeping the chip in a horizontal position (E) or by tilting it slightly (F) during spheroid growth. Scale bars: 50 µm* ***(G)*** *Fraction of wells containing a spheroid at day 7, for a given number of DLD-1 cells in the well at seeding.*

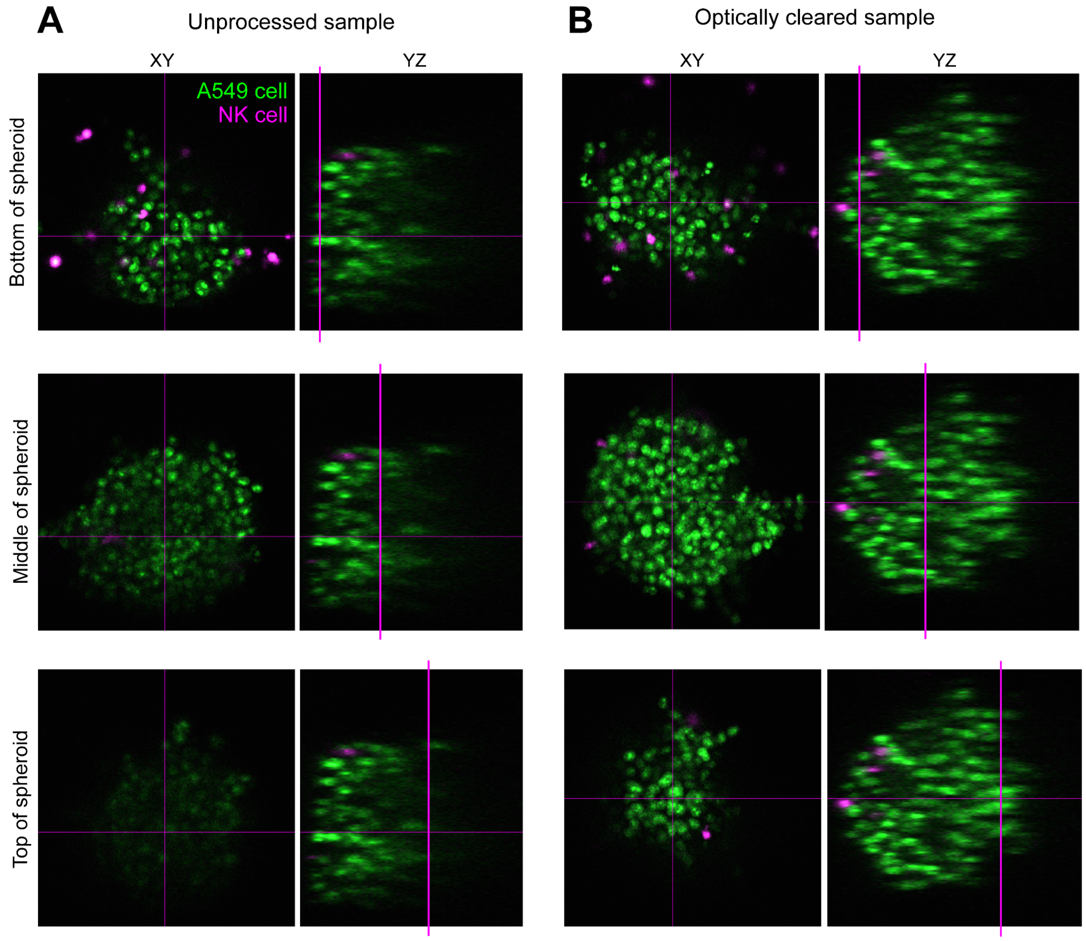


***Supporting figure S6. Improvement to deep imaging of spheroids by optical clearing.***
***(A-B)*** *Co-cultures of A549 spheroids and NK cells were fixed after 16 hours and either directly imaged with confocal microscopy (A), or first, permeabilized and embedded in a refractive index-matched medium before imaging (optical clearing) (B). The magenta cross indicates the cutting planes for the XY and YZ images.*

*Supporting Table 1. Spheroid formation in microwells using a range of immortal cell lines.*

| Cell line | Tissue | Cell type | Makes spheroids |
|---|---|---|---|
| DLD-1 | Colon | Epithelial | yes |
| HCT116 | Colon | Epithelial | yes |
| SW620 | Colon | Epithelial | yes |
| IMR-90 | Lung | Fibroblast | yes |
| MRC-5 | Lung | Fibroblast | yes |
| WI-38 | Lung | Fibroblast | yes |
| A549 | Lung | Epithelial | yes |
| A-498 | Kidney | Epithelial | yes |
| ACHN | Kidney | Epithelial | no |
| HeLa | Cervix | Epithelial | yes |
| IMR-32 | Brain | Neuroblast | yes |
| OVCAR-8 | Ovary | Epithelial | yes |
| MCF-7 | Breast | Epithelial | yes |
| BJ | Skin | Fibroblast | yes |
